# U2AF1 mutations rescue deleterious exon skipping induced by KRAS mutations

**DOI:** 10.1101/2025.03.21.644128

**Authors:** David M. Walter, Katherine Cho, Smruthy Sivakumar, Daniel Denney, Iris T.-H. Lee, Anders B. Dohlman, Jakob M. Heinz, Ethan Shurberg, Kevin X. Jiang, Akansha A. Gupta, Garrett M. Frampton, Matthew Meyerson

**Affiliations:** Department of Medical Oncology, Dana-Farber Cancer Institute, Boston, MA; Cancer Program, Broad Institute of Harvard and MIT, Cambridge, MA; Department of Genetics, Harvard Medical School, Boston, MA; Foundation Medicine Inc., Boston, MA; Department of Biomedical Informatics, Harvard Medical School, Boston, MA

## Abstract

The mechanisms by which somatic mutations of splicing factors, such as U2AF1^S34F^ in lung adenocarcinoma, contribute to cancer pathogenesis are not well understood. Here, we used prime editing to modify the endogenous *U2AF1* gene in lung adenocarcinoma cells and assessed the resulting impact on alternative splicing. These analyses identified *KRAS* as a key target modulated by U2AF1^S34F^. One specific *KRAS* mutation, G12S, generates a cryptic U2AF1 binding site that leads to skipping of *KRAS* exon 2 and generation of a non-functional *KRAS* transcript. Expression of the U2AF1^S34F^ mutant reverts this exon skipping and restores KRAS function. Analysis of cancer genomes reveals that U2AF1^S34F^ mutations are enriched in KRAS^G12S^-mutant lung adenocarcinomas. A comprehensive analysis of splicing factor/oncogene mutation co-occurrence in cancer genomes also revealed significant co-enrichment of KRAS^Q61R^ and U2AF1^I24T^ mutations. Experimentally, KRAS^Q61R^ mutation leads to *KRAS* exon 3 skipping, which in turn can be rescued by the expression of U2AF1^I24T^. Our findings provide evidence that splicing factor mutations can rescue splicing defects caused by oncogenic mutations. More broadly, they demonstrate a dynamic process of cascading selection where mutational events are positively selected in cancer genomes as a consequence of earlier mutations.

## Main

Analysis of recurrent somatic mutations in cancers through next-generation sequencing has identified novel classes of cancer mutated genes, including genes encoding components of the splicing machinery^1^. The splicing factor U2AF1 binds to an AG dinucleotide at the 3’ splice site of AG-dependent introns and recruits the U2 snRNP to these sites^2–4^. Mutations in *U2AF1* are found in acute myeloid leukemia (AML) and myelodysplastic syndrome (MDS)^1^, lung adenocarcinoma (LUAD)^5^ and pancreatic ductal adenocarcinoma (PDAC)^6^, and alter its affinity for AG dinucleotides depending on their nucleotide context, thereby disrupting normal splicing^7–10^.

U2AF1^S34F^ is the most common amino acid substitution mutation in LUAD after those in *KRAS* and *EGFR* (Extended Data Fig. 1a), suggesting a powerful selective force for this substitution^11^. The frequency of U2AF1^S34F^ mutations does not correlate with tobacco smoking frequency across cancer types (Extended Data Fig. 1b), and does not appear to correlate with APOBEC activity or with mutation rate based on sequence context^11–14^.

The mutation spectrum of *U2AF1* is distinct between AML/MDS, LUAD and PDAC. In AML/MDS, *U2AF1* mutations occur at one of two major hotspots, S34 or Q157 (Extended Data Fig. 1c)^11^. In contrast, there is a single dominant hotspot of *U2AF1* mutations in LUAD at S34, with 99% of these mutations being S34F (Extended Data Fig. 1c)^11,15^. Furthermore, in PDAC, while S34 is the primary hotspot, there is a secondary mutational hotspot at I24^11^. These mutational patterns suggest that there are distinct positive selective pressures for U2AF1 mutations across different cancer types.

Despite the frequency of *U2AF1* mutations in LUAD and PDAC, the functional impact of these mutations on cancer cell behavior is poorly understood. The majority of work on *U2AF1* mutant function has been performed in the context of AML and MDS, hematological cancers with very different mutational and transcriptomic contexts^7,11,16–18^. Investigations on the functions of U2AF1 mutations in carcinomas have led to multiple proposed mechanisms, including the regulation of epithelial-mesenchymal transition (EMT), mRNA translation, MAPK signaling and stress granule formation^7,15,18,19^. However, the specific selective advantages conferred by U2AF1^S34F^ and U2AF1^I24T^ mutations remain poorly understood.

To better delineate the function of *U2AF1* mutants, we combined prime editing of the endogenous locus with genomic analysis of cancer-derived mutations to uncover the selective pressure behind these mutations^20^.

## Generation of *U2AF1*-mutant lines by prime editing leads to differential alternative splicing

To identify the unique characteristics of U2AF1^S34F^ mutations that result in their selection in lung cancer, we applied twin prime editing (TwinPE)^20,21^. This allowed us to model a range of mutations at the S34 codon of endogenous *U2AF1* in LUAD cell lines and compare their impacts on alternative splicing to identify the unique characteristics of U2AF1^S34F^. As homozygous U2AF1 mutations are lethal to cells^22,23^, we used TwinPE to generate 6 heterozygous U2AF1 mutations in A549 cells. These consisted of a synonymous S34S mutation, S34C, S34A, S34Y, S34F and S34F(TTC) using an alternative codon for phenylalanine (Extended Data Table 1).

We performed RNA-sequencing on parental A549 cells as well as those modified at S34 (Extended Data Table 2). We then analyzed the alternative splicing events associated with each amino acid substitution and found that these were highest in cells harboring U2AF1^S34F^ mutations (Extended Data Fig. 2a). The expression of the prime edited alleles for each engineered *U2AF1* variant was similar except for a slight reduction in expression of U2AF1^S34F(TTC)^ (Extended Data Fig. 2b), associated with a slightly decreased splicing impact compared to U2AF1^S34F^ (Extended Data Fig. 2a). Alternative splicing events in U2AF1^S34F^- mutant A549 cells showed significant overlap with those from U2AF1^S34F^-mutant human LUAD samples^24^ (Extended Data Fig. 2c). In addition, the number of splicing events associated with each mutation positively correlated with the frequency of that mutation across human cancers (Extended Data Fig. 2d)^11,25,26^. U2AF1^S34F^ and U2AF1^S34F(TTC)^ mutations resulted primarily in skipped exons and alternative 3’ splice site usage (Extended Data Fig. 2e), and led to usage of CAG as opposed to TAG trinucleotides at the 3’ splice site (Extended Data Fig. 3), as reported previously for this mutation^8,18^.

## KRAS^G12S^-mutant cancers undergo *KRAS* exon 2 skipping which can be reversed by U2AF1^S34F^ mutations

To identify splicing events responsible for the positive selection of U2AF1^S34F^ mutations in lung cancer, we examined U2AF1^S34F^ and U2AF1^S34F(TTC)^-specific events that occurred in known oncogenes and tumor suppressors^27,28^. We identified splicing alterations in multiple genes of interest including *AXL, KRAS* and *NCOR2* (Fig. 1a). Combining mutation data from GENIE^11,29^ with alternative splicing analysis, *KRAS* was notable due to its high mutation frequency in U2AF1^S34F^-mutant LUAD (Extended Data Fig. 4a).

**Fig. 1:**
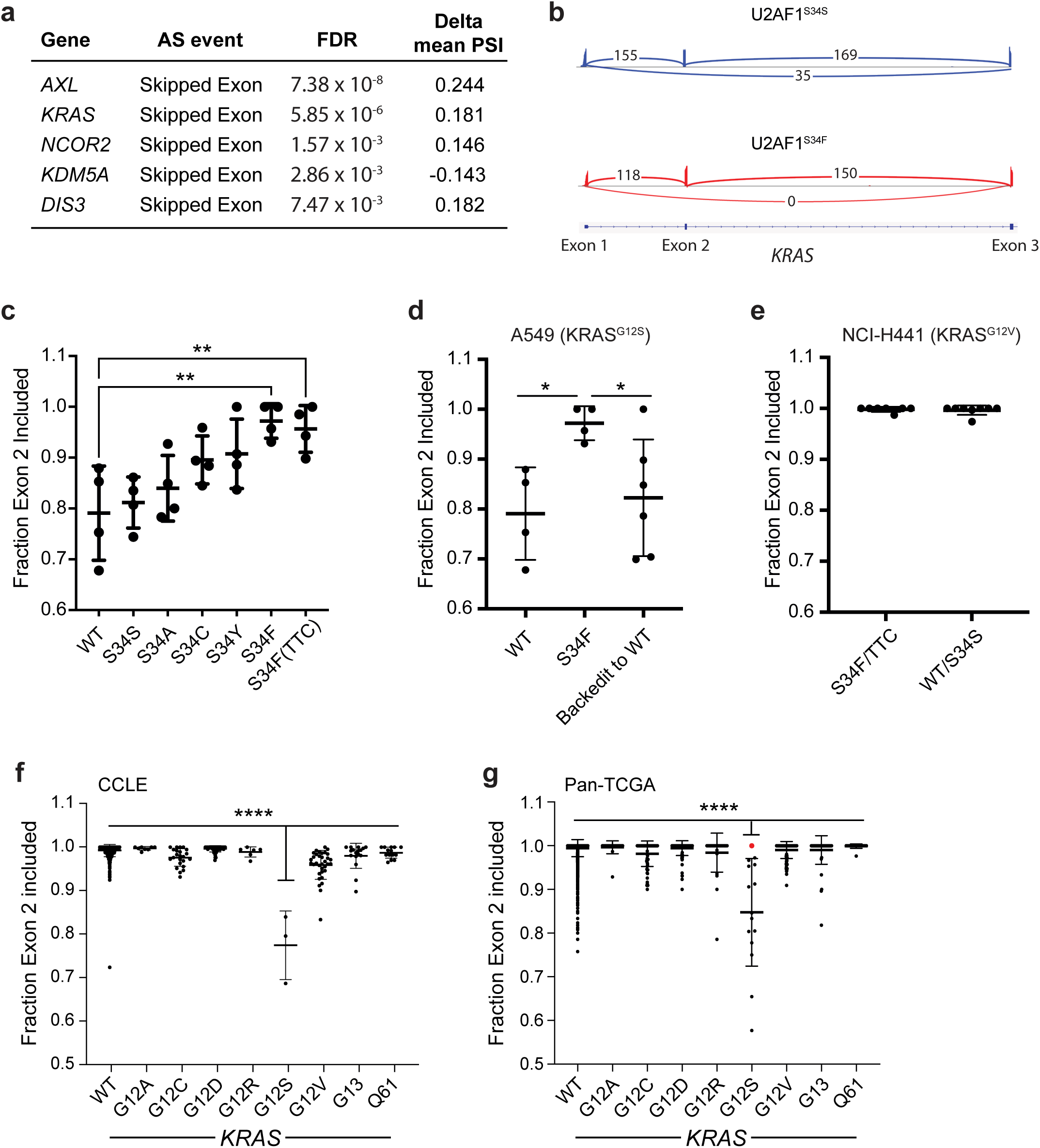
Cancers with KRAS^G12S^ mutations undergo *KRAS* exon 2 skipping which can be reversed by acquisition of U2AF1^S34F^ mutations. a) Table of the top alternative splicing events unique to U2AF1^S34F^ and U2AF1^S34F(TTC)^ -mutant cells in known oncogenes and tumor suppressors as defined by their presence in OncoKB^27,28^. Table shows the gene, type of alternative splicing event, false discovery rate-corrected p value, and difference in percent spliced in (PSI) between U2AF1^S34F^-mutant and parental A549 cells. b) Representative sashimi plots^80^ for the alternative splicing of *KRAS* exon 2 in A549 cells with U2AF1^S34S^ or U2AF1^S34F^ mutations. Sashimi plot was generated using Integrative Genomics Viewer^64^. c) Quantification of the fraction of RNA-sequencing reads with *KRAS* exon 2 inclusion across parental A549 cells or those harboring S34S, S34C, S34A, S34Y, S34F or S34F(TTC) mutations (n=4 clones for each). Significance shown for S34F (q ratio=4.25, DF=21, p=0.0019) and S34F(TTC) (q ratio=3.89, DF=21, p=0.0043). d) Quantification of fraction of RNA-sequencing reads with *KRAS* exon 2 inclusion for parental A549 cells (n= 4 clones), U2AF1^S34F^-mutant cells (n= 4 clones), or cells whose U2AF1^S34F^ mutations have been reverted to wildtype by prime editing (n= 6 clones). Statistical comparison between parental A549 cells and U2AF1^S34F^-mutant cells (t=3.67, df=6, 95% confidence interval=0.060 to 0.302, p=0.010) and between U2AF1^S34F^-mutant cells and backedited cells (t=2.44, df=8, 95% confidence interval= -0.291 to -0.0083, p=0.010) are shown. e) Quantification of the fraction of RNA-sequencing reads with *KRAS* exon 2 inclusion for NCI-H441 cells with U2AF1^S34F^ or U2AF1^S34F(TTC)^ mutations (n= 7 clones) or wildtype U2AF1 or U2AF1^S34S^ mutations (n=8 clones) (t=0.46, df=13, 95% confidence interval= -0.010 to 0.007, p=0.66). f) Quantification of the fraction of RNA-sequencing reads with *KRAS* exon 2 inclusion for cell lines from the Cancer Cell Line Encyclopedia (CCLE)^35^ with wildtype KRAS (n=835), KRAS^G12A^ (n=8), KRAS^G12C^ (n=23), KRAS^G12D^ (n=53), KRAS^G12R^ (n=6), KRAS^G12S^ (n=3), KRAS^G12V^ (n=34), KRAS^G13^ (n=17), or KRAS^Q61^ (n=14) mutations. Statistical analysis comparing KRAS^G12S^ to all other groups is shown (q ratio>19.27, DF=984, p=3.31x10^-13^ for all comparisons). g) Quantification of the fraction of RNA-sequencing reads with *KRAS* exon 2 inclusion for pan-cancer patient samples from The Cancer Genome Atlas (TCGA)^24^ with wildtype KRAS (n=5570), KRAS^G12A^ (n=24), KRAS^G12C^ (n=66), KRAS^G12D^ (n=120), KRAS^G12R^ (n=30), KRAS^G12S^ (n=15), KRAS^G12V^ (n=109), KRAS^G13^ (n=52), or KRAS^Q61^ (n=22) mutations. Statistical analysis comparing KRAS^G12S^ to all other groups is shown (q ratio>20.76, DF=5999, p=1.71x10^-12^ for all comparisons). Patient sample containing both a U2AF1^S34F^ mutation and concurrent KRAS^G12S^ mutation is marked in red.

*KRAS* is known to undergo alternative splicing to produce two dominant protein isoforms: KRAS4A and KRAS4B. The predicted protein isoforms differ in C-terminal regions that control membrane localization, with KRAS4A containing a palmitoylation domain and KRAS4B functioning via a poly-lysine sequence^30,31^, though both isoforms are capable of transforming cells^32^. In A549 cells with varying U2AF1 mutations, we found that *KRAS4A* was present in ∼5-12% of transcripts, in line with previous studies, but did not differ according to U2AF1 mutation status (Extended Data Fig. 4b).

Instead, we observed that exon 2 of *KRAS* was skipped in a median of 18% of reads (range 12-32%) in parental or U2AF1^S34S^-mutant A549 cells, while skipping was reduced to a median of 3% of reads (range 0-10%) in U2AF1^S34F^ and U2AF1^S34F(TTC)^-mutant cells (Fig. 1b,c, Extended Data Fig. 4c,d,e). Interestingly, exon 2 of *KRAS* harbors the protein’s translation start site as exon 1 is untranslated (Extended Data Fig. 4f). Exon 2 skipping is predicted to result in usage of a downstream alternative start codon in exon 3, omitting the first 66 amino acids of KRAS which include the P-loop, Switch-I and part of Switch-II (Extended Data Fig. 4g). Using long-read RNA-sequencing, we observed *KRAS* exon 2 skipping in both *KRAS4A* and *KRAS4B* transcripts, as well as in combination with previously described alternative 5’ and 3’UTRs^33^ (Extended Data Fig. 5).

To further probe the relationship between *KRAS* exon 2 skipping and U2AF1^S34F^, we “backedited” the S34F mutant allele to wildtype *U2AF1* and found that the frequency of the skipping event was restored to baseline levels (∼18% of reads) (Fig. 1d). To confirm this splicing event in another cell line, we edited the *U2AF1* locus in NCI-H441 cells. However, in contrast to A549 cells, NCI-H441 cells showed no exon 2 skipping regardless of *U2AF1* mutational status (Fig. 1e). As distinct *HRAS* mutations lead to *HRAS* exon 2 skipping or inclusion^34^, we examined the *KRAS* locus of the two cell lines, and noticed that while A549 cells harbor a KRAS^G12S^ mutation, NCI-H441 cells contain a KRAS^G12V^ mutation.

To better understand the link between *KRAS* mutation status and alternative splicing, we examined alternative splicing data from 993 cancer cell lines from the Cancer Cell Line Encyclopedia (CCLE)^35^. We found that only KRAS^G12S^-mutant cell lines such as A549 cells had appreciable levels of *KRAS* exon 2 skipping (Fig. 1f, Extended Data Fig. 6a). The same held true in 6008 cancers with reported *KRAS* splicing data from The Cancer Genome Atlas (TCGA)^24^, where *KRAS* exon 2 skipping was minimal except in cancers with KRAS^G12S^ mutations (Fig. 1g, Extended Data Fig. 6b). Furthermore, we found a negative correlation between variant allele frequency (VAF) of KRAS^G12S^ and exon 2 inclusion, suggesting that tumors with a higher fraction of KRAS^G12S^-mutant cells have a higher degree of exon 2 skipping (Extended Data Fig. 6c). The TCGA dataset also contained a KRAS^G12S^-mutant case which had 100% *KRAS* exon 2 inclusion (Fig. 1g, Extended Data Fig. 6c, red dot). Remarkably, this case contained both KRAS^G12S^ and U2AF1^S34F^ mutations, consistent with our findings that U2AF1^S34F^ restores exon 2 inclusion.

In addition to *KRAS* exon 2 skipping, we examined the relative usage of *KRAS4A* and *KRAS4B* across KRAS-mutant cancer cells and patient samples. In CCLE data, we found that most *KRAS* mutations trended towards increased *KRAS4A* usage (Extended Data Fig. 6d). We also investigated *KRAS4A* usage in 7518 cancers with *KRAS4A/KRAS4B* splicing data from TCGA and found that each assessed *KRAS* mutation was associated with significantly higher usage of *KRAS4A* than samples with wildtype KRAS (Extended Data Fig. 6e). These data support previous findings that *KRAS* mutations in LUAD are associated with increased levels of *KRAS4A* usage^36^.

## KRAS^G12S^-mutant RNA contains a cryptic U2AF1 binding site which leads to exon 2 skipping

We observed that the RNA sequence of the KRAS^G12S^ mutation resembles a U2AF1 binding site, creating a UAG trinucleotide predicted to be bound by wildtype U2AF1 protein but not U2AF1^S34F^ (Fig. 2a, Extended Data Fig. 7a)^8,37^. Previous work has reported that expression of KRAS^G12V^ or KRAS^Q61H^ leads to decreased phosphorylation of SR splicing factors and global changes in alternative splicing^38^. Therefore, we wondered if the *KRAS* exon 2 skipping we observed was due to the KRAS^G12S^ amino acid substitution itself, or due to the creation of a cryptic U2AF1 binding site in the RNA. To test this, we performed prime editing on A549 U2AF1^S34S^ cells to mutate the homozygous KRAS^G12S^ sequence to another codon for serine (TCT) that does not resemble a U2AF1 binding site (Fig. 2a). RT-PCR and RNA-sequencing analysis of exon 2 skipping found that exon skipping was completely abolished in KRAS^G12S^- mutant cells with a TCT codon (Fig. 2b,c). This demonstrates that the creation of a novel cryptic U2AF1 binding site is required for *KRAS* exon 2 skipping, as opposed to the G12S amino acid substitution itself.

**Fig. 2:**
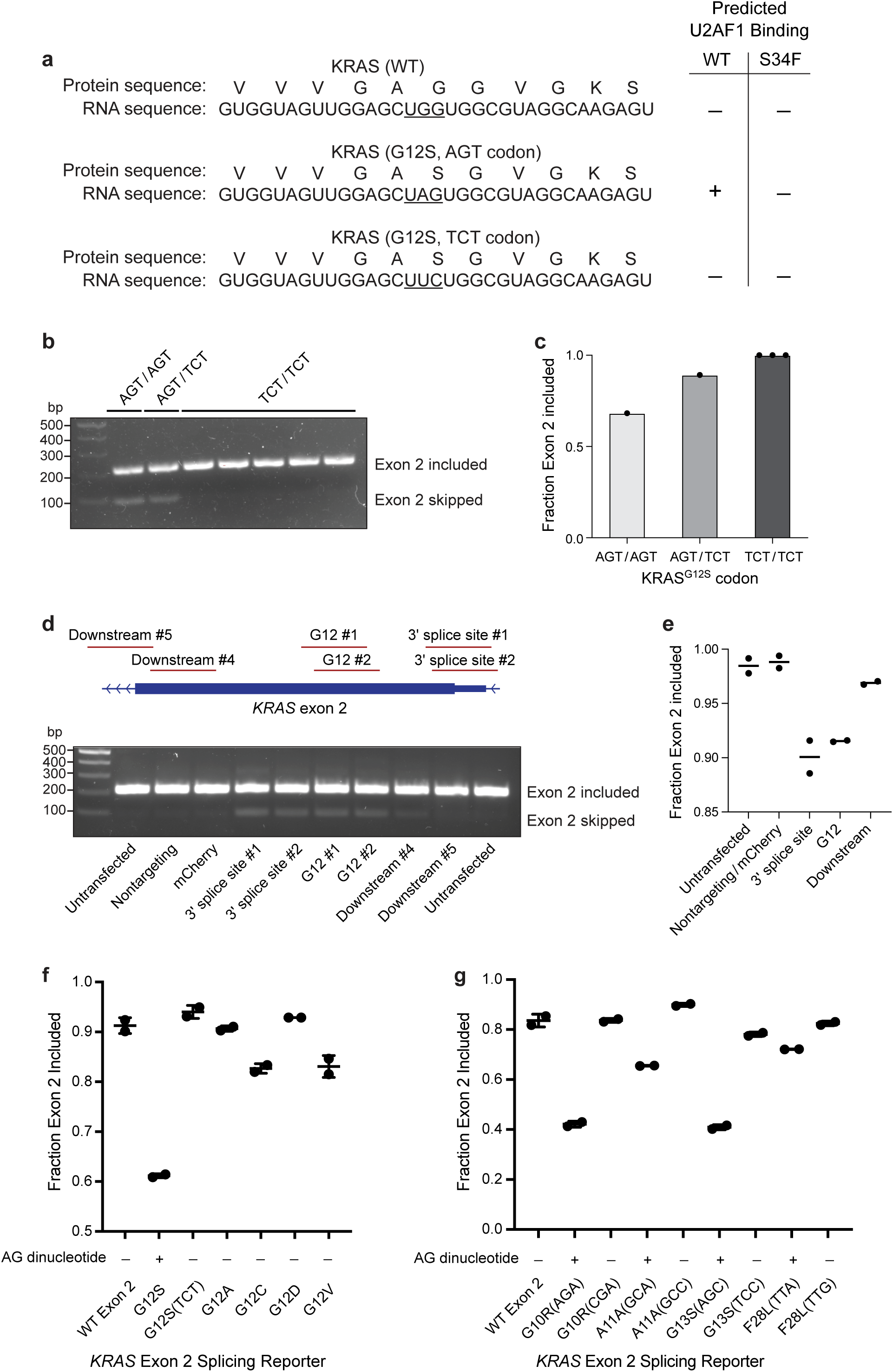
The RNA sequence encoding KRAS^G12S^ creates a cryptic U2AF1 binding site which disrupts normal exon 2 inclusion. a) RNA and protein sequence for wildtype KRAS and KRAS^G12S^ indicating a cryptic U2AF1 binding site which is predicted to be bound by wildtype U2AF1 but not U2AF1^S34F^. KRAS^G12S^ using an alternative codon for serine (TCT) does not resemble a U2AF1 binding site. b) RT-PCR detection of *KRAS* exon 2 skipping for A549 cells with KRAS^G12S^ mutations using the codons AGT/AGT (n=1 clone), AGT/TCT (n=1 clone) or TCT/TCT (n=3 clones). c) Quantification of the fraction of RNA-sequencing reads with *KRAS* exon 2 inclusion in A549 cells with AGT/AGT, AGT/TCT or TCT/TCT (n = 3 clones) codons. d) Schematic of dCasRx sgRNA locations and agarose gel for RT-PCR detection of *KRAS* exon 2 skipping in NCI-H2023 cells. Cells were transfected with dCasRx in addition to nontargeting sgRNA (n=1) or sgRNAs targeting mCherry RNA (n=1), the 3’ splice site of *KRAS* RNA (n=2), the G12 sequence of *KRAS* RNA (n=2), or downstream regions of *KRAS* RNA (n=2). e) Quantification of the fraction of *KRAS* exon 2 inclusion from d) in untransfected NCI-H2023 cells compared to cells following treatment with dCasRx alongside nontargeting or mCherry targeting sgRNA (n=2), 3’ splice site sgRNA (n=2), G12 sgRNA (n=2) or downstream sgRNA (n=2). f) Quantification of the fraction of *KRAS* Exon 2 inclusion using a *KRAS* exon 2 splicing reporter and measured by RT-PCR from Extended Data Fig. 7c. 293T cells were transfected in biological duplicate with splicing reporter DNA containing wildtype *KRAS* exon 2 (n=2), or *KRAS* exon 2 with common mutations including G12S (n=2), G12S(TCT) (n=2), G12A (n=2), G12C (n=2), G12D (n=2) or G12V (n=2). g) Quantification of the fraction of *KRAS* Exon 2 inclusion using a *KRAS* exon 2 splicing reporter and measured by RT-PCR from Extended Data Fig. 7d. 293T cells were transfected in biological duplicate with splicing reporter DNA containing wildtype *KRAS* exon 2 (n=2), or *KRAS* exon 2 with mutations introducing novel U2AF1 binding sites (AG dinucleotides) or synonymous mutations without U2AF1 binding sites. Mutations that generated novel AG dinucleotides include G10R(AGA) (n=2), A11A(GCA) (n=2), G13S(AGC) (n=2) and F28L(TTA) (n=2), while those that do not contain AG dinucleotides include G10R(CGA) (n=2), A11A(GCC) (n=2), G13S(TCC) (n=2), and F28L(TTG) (n=2).

As it seemed surprising that a cryptic U2AF1 binding site would cause exon skipping rather than alternative 3’ splice site usage, we performed a thorough analysis of the RNA-Seq reads using Integrative Genomics Viewer (IGV 2.16.0)^39^. Alternative 3’ splice site usage would correlate with a stepwise increase in read coverage near the G12 RNA sequence, similar to the 3’UTR of *CTNNB1*, where U2AF1^S34F^ leads to decreased usage of the distal 3’ splice site^8^ (Extended Data Fig. 7b). However, we observed relatively constant read coverage across *KRAS* exon 2 regardless of U2AF1 mutational status, confirming that KRAS^G12S^ leads to exon 2 skipping as opposed to alternative 3’ splice site usage (Extended Data Fig. 7b).

As the 3’ splice site upstream of *KRAS* exon 2 is only 45 bp from the cryptic binding site at G12S, we hypothesized that exon 2 skipping might arise from steric hindrance caused by wildtype U2AF1 binding and recruiting the spliceosome complex to this alternative site. To test whether such a process was possible, we targeted a catalytically inactive version of the RNA binding CasRX protein (dCasRx)^40^ to various locations spanning *KRAS* exon 2 in NCI-H2023 cells with wildtype *KRAS* (Extended Data Table 3). Indeed, binding of dCasRx to the canonical 3’ splice site leads to *KRAS* exon 2 skipping, likely due to interference with normal U2AF1 binding (Fig. 2d,e). Remarkably, directing dCasRx to the G12 sequence also resulted in significant levels of exon 2 skipping, while binding ∼60 bases downstream lessened the impact on exon 2 skipping, likely due to decreased steric hindrance further from the upstream 3’ splice site (Fig. 2d,e). This suggests that aberrant U2AF1 binding to the G12S codon could lead to *KRAS* exon 2 skipping due to steric hindrance with the proximal 3’ splice site.

To directly assess the impact of *KRAS* sequence variation on splicing, we chose to use a simplified genomic system: a splicing reporter minigene using the pSpliceExpress system^41,42^. Here we incorporated exon 2 of *KRAS* and 500 bp of each flanking intron between 2 constitutively expressed exons of the rat insulin gene. We found that the introduction of G12S mutations resulted in significant skipping of *KRAS* exon 2, while other mutations at G12 had minimal to no effect, demonstrating that KRAS^G12S^ mutations directly lead to exon skipping (Fig. 2f, Extended Data Fig. 7c). As observed in our edited A549 cells, KRAS^G12S^ using a TCT codon resulted in no exon skipping (Fig. 2f, Extended Data Fig. 7c).

One mechanistic hypothesis for *KRAS* exon 2 skipping driven by KRAS^G12S^ mutation is the introduction of a novel U2AF1 binding site containing an AG dinucleotide. If this hypothesis is correct, then introducing other AG-containing U2AF1 binding sites could similarly cause exon skipping. Using our splicing reporter minigene, we generated a series of nucleotide substitutions upstream of or near the G12 codon of *KRAS*. We observed significant levels of *KRAS* exon 2 skipping for each variant that introduces an AG dinucleotide and less exon 2 skipping for any variant that did not introduce an AG dinucleotide (Fig. 2g, Extended Data Fig. 7d). Additionally, exon skipping was less pronounced for a U2AF1 binding site created further downstream from the 3’SS (Fig. 2g, Extended Data Fig. 7d). Taken together, these results suggest that KRAS^G12S^ mutation promotes exon skipping via the creation of a new AG-containing U2AF1 binding site.

## *KRAS* exon 2 skipping leads to reduced MAPK signaling and cell growth

While the effects of *KRAS* exon 2 skipping have not been studied, a similar event has been identified in *HRAS* whereby exon 2 skipping is thought to lead to decreased signaling in the context of Costello syndrome^34^. Therefore, we hypothesized that *KRAS* exon 2 skipping might lead to decreased MAPK signaling, while acquisition of U2AF1^S34F^ mutations would enhance KRAS signaling by restoring exon 2 inclusion. We used flow cytometry to measure phosphorylation of Erk as a readout for MAPK signaling and found that that the acquisition of U2AF1^S34F^ mutations increased p-Erk levels by ∼37% in KRAS^G12S^-mutant A549 cells (Fig. 3a,b). Similarly, A549 cells in which the homozygous KRAS^G12S^ mutation was mutated to a TCT codon had a ∼64% increase in p-Erk levels (Extended Data Fig. 8a,b). The higher levels of p-Erk in cells harboring the KRAS^G12S(TCT)^ mutation compared to those with U2AF1^S34F^ may be in part due to “locking in” *KRAS* to a state of 100% exon 2 inclusion through the removal of the cryptic U2AF1 binding site.

**Fig. 3:**
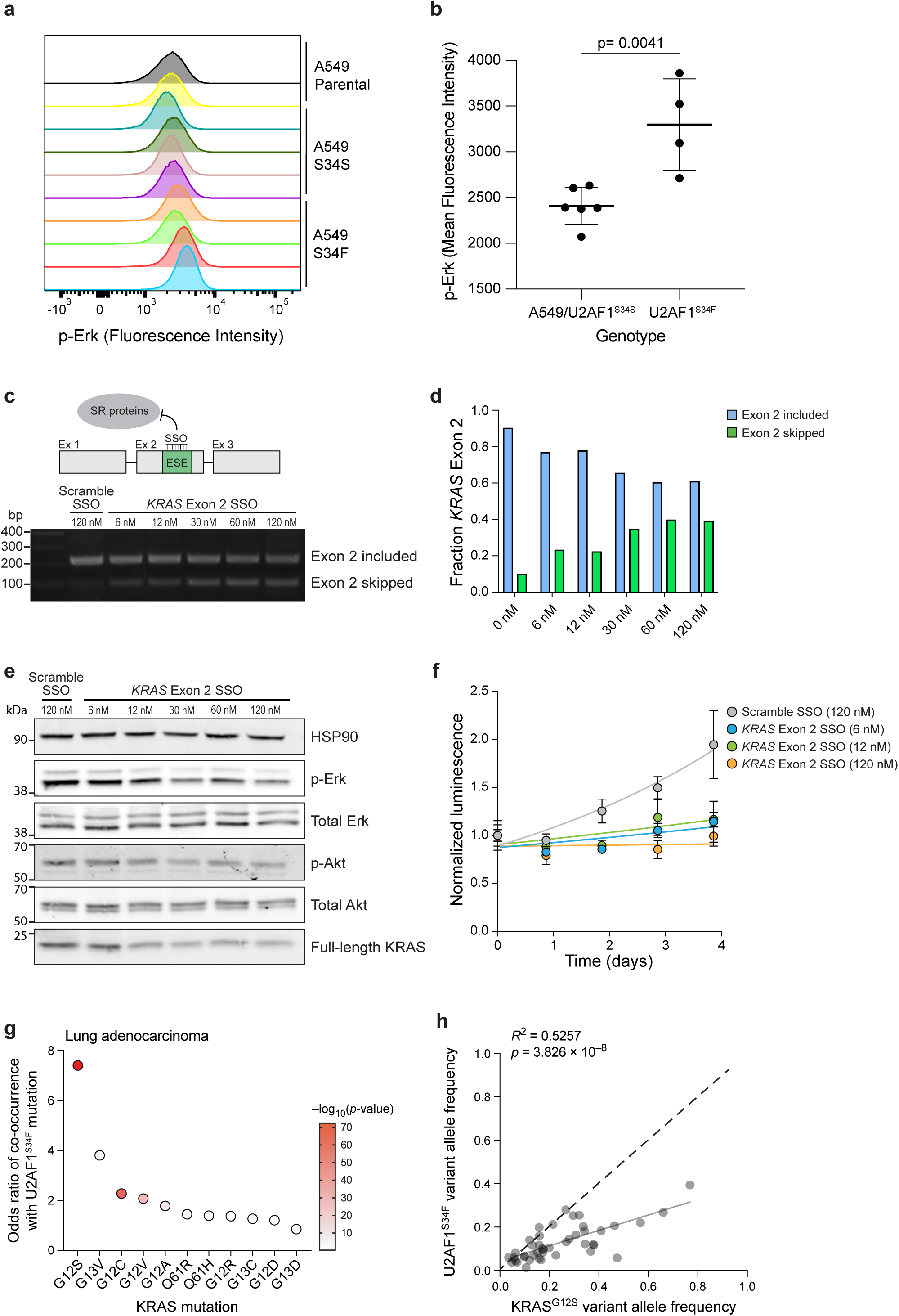
*KRAS* exon 2 skipping limits MAPK signaling and cell growth, leading to the positive selection of U2AF1^S34F^ mutations in KRAS^G12S^-mutant lung adenocarcinomas. a) Fluorescence intensity of p-Erk as measured by flow cytometry for parental A549 cells (n= 2 clones), U2AF1^S34S^-mutant A549 cells (n= 4 clones), or U2AF1^S34F^-mutant A549 cells (n= 4 clones). b) Quantification of the mean fluorescence intensity of p-Erk staining in parental A549 or U2AF1^S34S^ cells (n= 6 clones) or U2AF1^S34F^ cells (n= 4 clones) (t=3.98, df=8, 95% confidence interval = 373 to 1401, p=0.0041). c) Schematic indicates the binding of splice switching oligonucleotides (SSOs) to exonic splicing enhancers (ESEs) in *KRAS* exon 2, blocking SR protein binding and inhibiting exon 2 inclusion. RT-PCR gel demonstrates the detection of *KRAS* exon 2 skipping for NCI-H23 cells treated with scramble SSO or increasing concentrations of *KRAS* exon 2 SSO. d) Quantification of the fraction of transcripts with *KRAS* exon 2 skipping or inclusion from c). e) Immunoblot analysis of HSP90, p-Erk, total Erk, p-Akt, total Akt, and full length KRAS protein in NCI-H23 cells treated with increasing concentrations of *KRAS* exon 2 skipping SSOs 3 days after treatment. f) Cell viability of NCI-H23 cells grown on ultra-low attachment plates and treated with increasing concentrations of *KRAS* exon 2 skipping SSOs over a period of 5 days as measured by CellTiter-Glo (n= 4 technical replicates at each time point). g) Odds ratio of co-occurrence for U2AF1^S34F^ mutations and indicated KRAS mutations in lung adenocarcinoma patients using data from Foundation Medicine Inc. (n=62009 patients) and AACR Project GENIE (v14.0, n=14908 patients)^11^. h) Correlation between the variant allele frequencies for KRAS^G12S^ and U2AF1^S34F^ in 43 patient samples from the Foundation Medicine Inc. dataset containing both mutations and lacking CNVs at either locus (F=45.4, DFn=1, DFd=41, R^2^=0.526, p=3.83x10^-8^). Dashed line indicates a 1:1 ratio of KRAS^G12S^ and U2AF1^S34F^ allele frequencies.

To further test the impact of *KRAS* exon 2 inclusion on cell growth and signaling, we employed splice switching oligonucleotides (SSOs) targeting *KRAS* exon 2 (Extended Data Table 4)^34,43^. Treatment of NCI-H23 cells (KRAS^G12C^) with increasing concentrations of exon-skipping SSOs led to increased levels of exon 2 skipping (Fig. 3c,d). While canonical KRAS signaling proceeds through a RAF/MEK/ERK axis, there is evidence that KRAS-mediated PI3K/AKT/mTOR signaling is also important for tumor initiation and maintenance^44^. We found that higher levels of *KRAS* exon 2 skipping led to decreased signaling through both pathways as indicated by reduced levels of p-Erk and p-Akt, as well as full length KRAS protein (Fig. 3e). We also treated cells with increasing levels of exon-skipping SSOs and grew them on ultra-low attachment plates, where we observed even small increases in *KRAS* exon 2 skipping led to decreased cell growth (Fig. 3f). These findings indicate that *KRAS* exon 2 skipping limits KRAS signaling and is detrimental to cell proliferation.

## There is positive selection for U2AF1^S34F^ mutations in KRAS^G12S^-mutant cancers

Our data show that KRAS^G12S^ mutations result in exon 2 skipping which can be reversed by U2AF1^S34F^ mutations, and that exon 2 skipping leads to reduced MAPK signaling and cell growth. Therefore, we hypothesized that U2AF1^S34F^ mutations would be favored in KRAS^G12S^- mutant cancers as a means of driving KRAS signaling and tumor growth. To determine whether this was the case, we examined a set of 76,917 LUAD cases including 62,009 from Foundation Medicine and 14,908 from AACR Project GENIE and calculated the odds ratio of co-occurrence between U2AF1^S34F^ and varying *KRAS* mutations^11,26^. Remarkably, we found that KRAS^G12S^ mutation had an odds ratio of 7.4 for U2AF1^S34F^ mutations, far higher than any other *KRAS* mutation, with a p value of 4.8 x 10^-73^ (Fig. 3g, Extended Data Fig. 8c). Interestingly, KRAS^G12C^ and KRAS^G12V^ mutations were also significantly enriched, though to a lesser degree, and displayed intermediate levels of exon 2 skipping in our splicing reporter assay. In addition, we found significant co-occurrence of U2AF1^S34F^ mutations and KRAS^G12S^ mutations in pancreatic adenocarcinoma cases (Extended Data Fig. 8d).

U2AF1^S34F^ has been described as a truncal mutation in both hematological malignancies and LUAD^15,45,46^. However, as U2AF1^S34F^ rescues cryptic exon 2 skipping in KRAS^G12S^-mutant cancers, we could imagine that U2AF1^S34F^ mutations might occur as secondary events after the acquisition of a KRAS^G12S^ mutation. To test this possibility, we analyzed the variant allele frequencies (VAF) of KRAS^G12S^ and U2AF1^S34F^ in 43 LUAD cases from the Foundation Medicine Inc. dataset which harbored both mutations and lacked copy number variations (CNVs) at either locus. We found a strong positive correlation between the two mutations, however the VAF for KRAS^G12S^ was significantly higher than the VAF for U2AF1^S34F^ across the tumors, suggesting that U2AF1^S34F^ occurs as a secondary mutation in KRAS^G12S^-mutant LUADs (Fig. 3h, Extended Data Fig. 8e).

Spliceosome inhibitors have gained recent interest as potential clinical targets, with a particular focus on compounds targeting the splicing factor SF3B1. It is possible that inhibition of SF3B1 may disrupt normal splicing of *KRAS* and limit MAPK signaling. To test this possibility, we treated NCI-H2023 cells with increasing concentrations of SF3B1 inhibitor Pladienolide B^47,48^ and tested its impact on *KRAS* exon 2 skipping and MAPK signaling as measured by phosphorylation of Erk. We found that treatment with Pladienolide B led to increased *KRAS* exon 2 skipping (Extended Data Fig. 8f) and decreased p-Erk after 20 hours of treatment (Extended Data Fig. 8g,h).

## KRAS^Q61R/Q61L^ mutations lead to *KRAS* exon 3 skipping and are rescued by U2AF1^I24T^ mutation

We searched for additional splicing factor/oncogene relationships by examining mutations which caused exon skipping events within the mutated gene and which had a significant co-occurrence with mutations in the splicing factor genes *U2AF1*, *SF3B1* or *SRSF2* (Fig. 4a). Interestingly, the strongest relationship was between another pair of *KRAS* and *U2AF1* mutations: KRAS^Q61R^ and U2AF1^I24T^ (Fig. 4a).

**Fig. 4:**
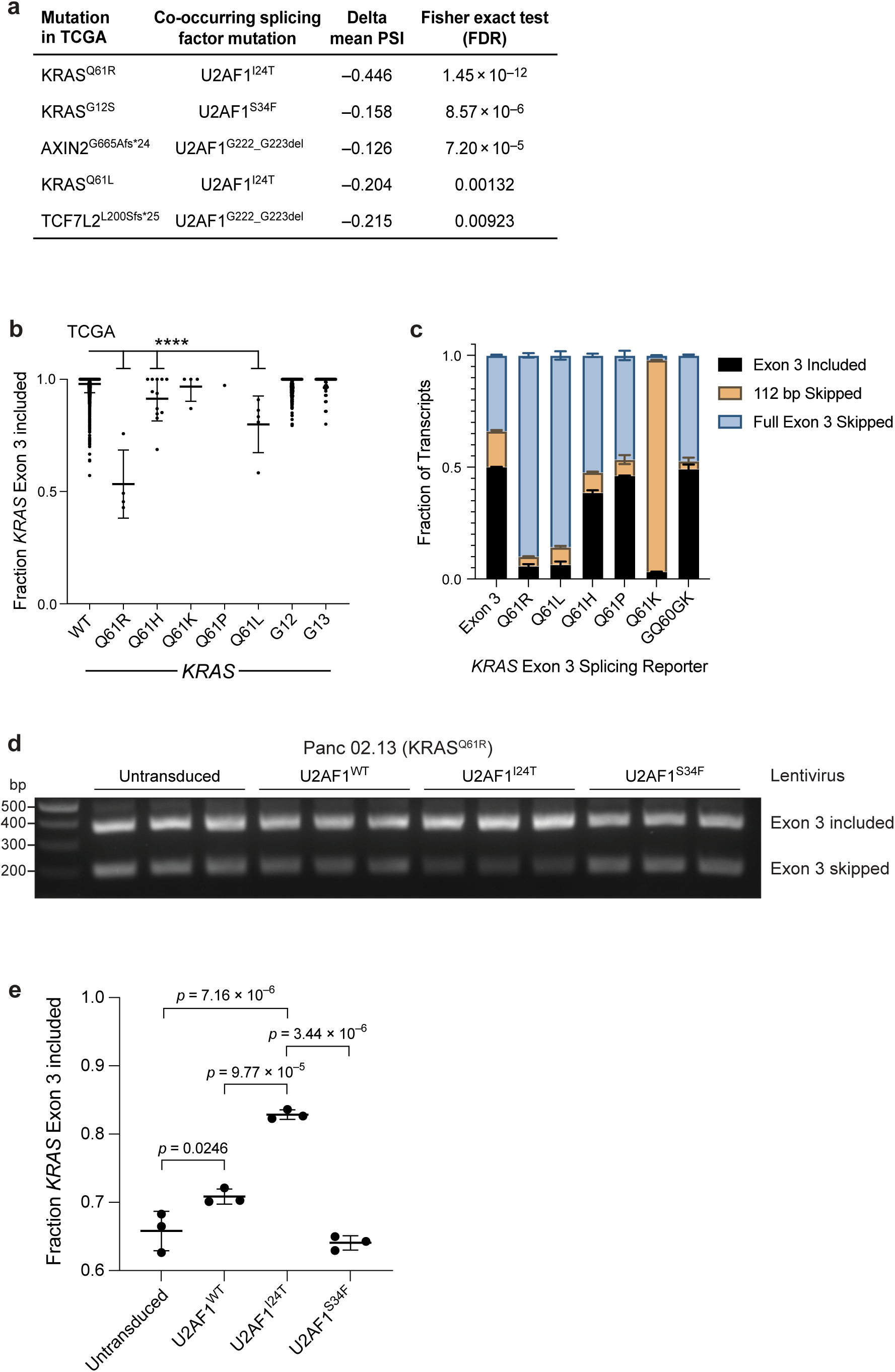
KRAS^Q61R/L^-mutant pancreatic cancers acquire secondary U2AF1^I24T^ mutations that rescue inadvertent skipping of *KRAS* exon 3. a) Table of mutations associated with exon skipping events in TCGA data^24^, and which co-occur with splicing factor mutations (data from AACR Project GENIE^11^). Threshold for exon skipping: delta mean PSI <-0.1. Threshold for co-occurrence: p value < 0.00001 as determined by Fisher’s exact test. b) Quantification of the fraction of RNA-sequencing reads with *KRAS* exon 3 inclusion for pan-cancer patient samples from TCGA^24^ with wildtype KRAS (n=6065), KRAS^Q61R^ (n=4), KRAS^Q61H^ (n=12), KRAS^Q61K^ (n=4), KRAS^Q61P^ (n=1), KRAS^Q61L^ (n=5), KRAS^G12^ (n=388), or KRAS^G13^ (n=53) mutations. Statistical comparisons between wildtype KRAS and KRAS^Q61R^ (q ratio=22.3, DF=6524, p<1x10^-15^), KRAS^Q61H^ (q ratio=5.71, DF=6524, p=8.2x10^-8^), and KRAS^Q61L^ (q ratio=10.08, DF=6524, p<1x10^-15^) are shown. c) Quantification of the fraction of *KRAS* Exon 3 inclusion, skipping of a 112 bp fragment of exon 3, or skipping the entirety of exon 3 using a *KRAS* exon 3 splicing reporter. Splicing was measured by RT-PCR from Extended Data Fig. 9b. 293T cells were transfected in biological duplicate with splicing reporter DNA containing wildtype *KRAS* exon 3 (n=2), or *KRAS* exon 3 with common mutations including Q61R (n=2), Q61L (n=2), Q61H (n=2), Q61P (n=2), Q61K (n=2) or GQ60GK (n=2) mutation. d) RT-PCR detection of *KRAS* exon 3 skipping for untransduced KRAS^Q61R^-mutant Panc 02.13 cells (n= 3 biological replicates) or after delivery of lentivirus expressing wildtype U2AF1 (n=3 biological replicates), U2AF1^I24T^ (n=3 biological replicates) or U2AF1^S34F^ (n=3 biological replicates). e) Quantification of the fraction of *KRAS* exon 3 skipping in Fig. 4d. Statistical comparisons for untransduced vs. U2AF1^WT^ (q ratio=5.252, DF=8, p=0.0246), untransduced vs. U2AF1^I24T^ (q ratio=17.76, DF=8, p=7.16x10^-6^), U2AF1^WT^ vs. U2AF1^I24T^ (q ratio=12.51, DF=8, p=9.77x10^-5^), and U2AF1^I24T^ vs. U2AF1^S34F^ (q ratio=19.57, DF=8, p=3.44x10^-6^) are shown.

Our computational analysis suggested that KRAS^Q61R^ mutations lead to significant levels of *KRAS* exon 3 skipping. Like *KRAS* exon 2, exon 3 skipping is expected to result in non-functional KRAS due to a frameshift after the first 37 amino acids of the protein, resulting in loss of Switch-II and all subsequent domains (Extended Data Fig. 9a). To understand the impact of KRAS^Q61R^ mutations on *KRAS* splicing more clearly, we examined exon 3 skipping in 6,532 TCGA samples with *KRAS* exon 3 skipping data^24^. Unlike KRAS mutations at G12, where only G12S resulted in significant exon skipping, multiple mutations at Q61 led to *KRAS* exon 3 skipping, including Q61R, Q61L and Q61H (Fig. 4b). Interestingly, work from Kobayashi *et al*. found that KRAS^Q61K^ mutations lead to either complete exon 3 skipping or skipping of 112 bp of the exon^49^. However, this is rescued by the acquisition of adjacent silent mutations, explaining the absence of exon 3 skipping in Q61K-mutant samples.

To further interrogate the impact of Q61 mutations on *KRAS* exon 3 inclusion, we again employed the pSpliceExpress splicing reporter^41,42^. By incorporating exon 3 of *KRAS* and 500 bp each of flanking intronic sequence into the vector, we found that Q61R and Q61L mutations led to dramatic exon 3 skipping, while Q61P and Q61H mutations resembled wildtype Exon 3 (Fig. 4c and Extended Data Fig. 9b). Interestingly, when we introduced a Q61K mutation, close to 100% of transcripts skipped 112 bp of exon 3, a splicing event previously identified by Kobayashi *et al.*^49^. However, as described^49^, a silent mutation in G60 restored exon 3 inclusion to similar levels as the wildtype reporter.

I24 of U2AF1 is located between its first and second RNA-binding zinc finger domains^50^, consistent with an impact on RNA binding specificity. Indeed, a study of rare spliceosomal mutations found that U2AF1^I24T^ leads to preferential inclusion of exons containing CAGG sequences at the 3’ splice site^51^. This CAGG motif is the exact sequence of the 3’ splice site of *KRAS* exon 3, suggesting that the acquisition of U2AF1^I24T^ mutations may restore inclusion of *KRAS* exon 3 in cells with KRAS^Q61^ mutations. To test this concept, we attempted to mutate the endogenous *U2AF1* locus using prime editing, however this was not successful. Therefore, we cloned wildtype U2AF1, U2AF1^S34F^ and U2AF1^I24T^ into the pLX301 lentiviral backbone^52^, and transduced the resulting vectors into KRAS^Q61R^-mutant Panc 02.13 cancer cells. Remarkably, we found that expressing U2AF1^I24T^, but not U2AF1^S34F^, rescued *KRAS* exon 3 inclusion from ∼66% of transcripts to ∼83% of transcripts (Fig. 4d,e). Interestingly, we also observed that over-expression of wildtype U2AF1 resulted in a small but statistically significant increase in *KRAS* exon 3 inclusion (Fig. 4d,e).

## *KRAS* exon 3 skipping is detrimental to cell growth, while the acquisition of secondary U2AF1^I24T^ mutations is selected for in KRAS^Q61R/L^-mutant cases

We previously observed that *KRAS* exon 2 skipping restricts cell growth and hypothesized that the same may be true for exon 3. To test whether *KRAS* exon 3 skipping hindered cell growth as expected, we employed SSOs targeting exon 3. As before, delivery of increasing amounts of exon 3 SSO led to increased exon skipping as quantified by RT-PCR (Fig. 5a,b). Furthermore, we found that KRAS^G12C^-mutant NCI-H23 cells were sensitive to *KRAS* exon 3 skipping, with relatively small increases in skipping leading to significant reductions in cell growth (Fig. 5c).

**Fig. 5:**
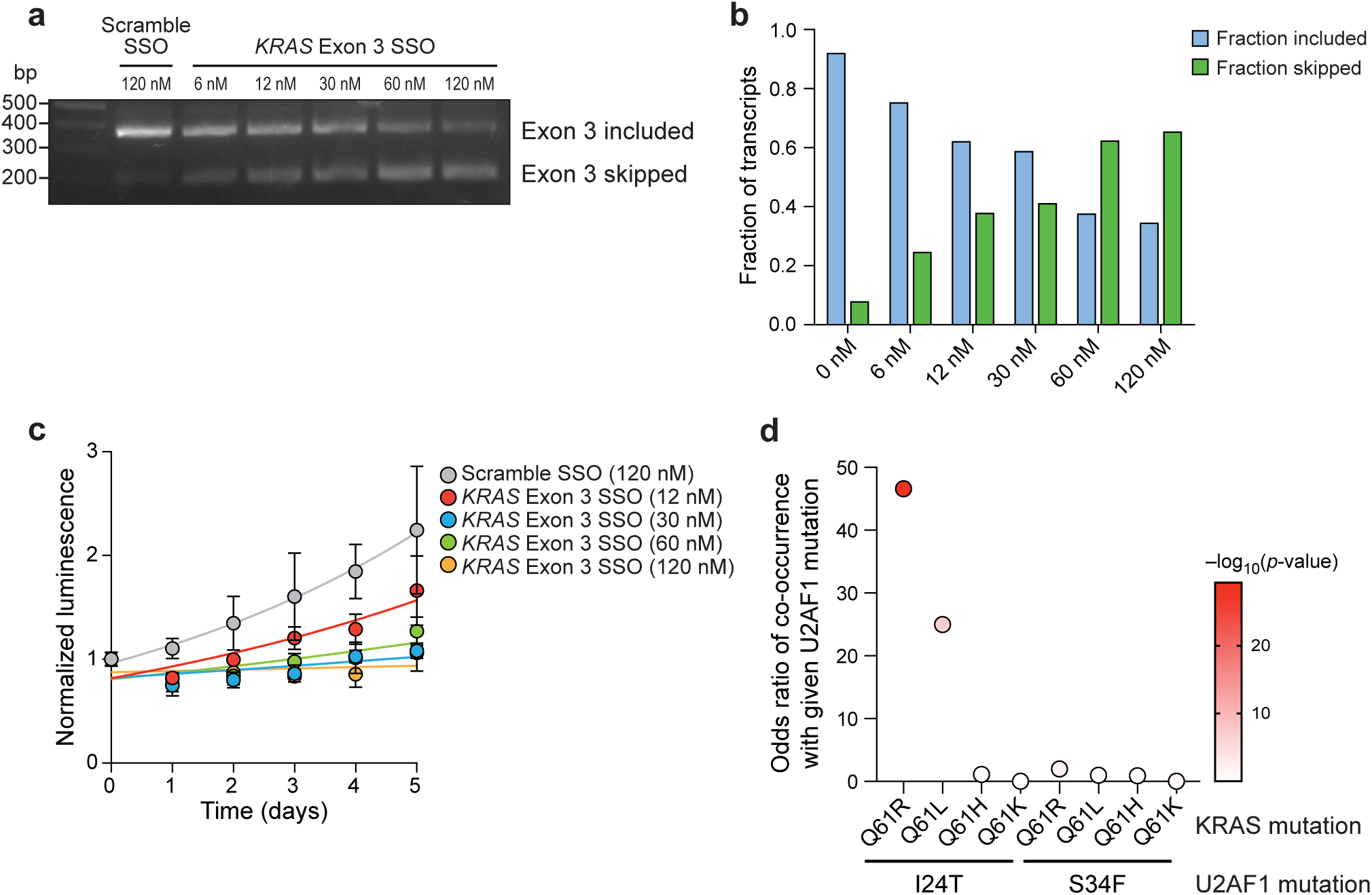
*KRAS* exon 3 skipping leads to reduced cell growth, while U2AF1^I24T^ mutations are enriched in KRAS^Q61R/L^-mutant pancreatic cancers. a) RT-PCR detection of *KRAS* exon 3 skipping for NCI-H23 cells treated with scramble SSO or increasing concentrations of *KRAS* exon 3 SSO. b) Quantification of the fraction of transcripts with *KRAS* exon 3 skipping or inclusion from a). c) Cell viability of NCI-H23 cells grown on ultra-low attachment plates and treated with increasing concentrations of *KRAS* exon 3 skipping SSOs over a period of 5 days as measured by CellTiter-Glo (n= 4 technical replicates at each time point). d) Odds ratio of co-occurrence for U2AF1^I24T^ or U2AF1^S34F^ mutations with KRAS^Q61R^, KRAS^Q61L^, KRAS^Q61H^ or KRAS^Q61K^ mutations in pancreatic cancers using data from Foundation Medicine Inc. (n=31,530 patients) and AACR Project GENIE (v16.1, n=8,304 patients)^11^.

As a result, we would expect enrichment of secondary U2AF1^I24T^ mutations in KRAS^Q61R/L^-mutant cancers. Patient data from AACR Project GENIE reveals that U2AF1^I24T^ mutations are ∼4 times more likely to occur in pancreatic cancer compared to other cancer types^11^. Therefore, we interrogated genomic data of 31,530 pancreatic cancer cases from Foundation Medicine Inc. and 8,304 cases from AACR Project GENIE (v16.1) to determine the odds ratio of co-occurrence for U2AF1^I24T^ and various mutations at Q61. Remarkably, U2AF1^I24T^ and KRAS^Q61R^ had an odds ratio for co-occurrence of 46.6, while an odds ratio of 25 was calculated for the co-occurrence of U2AF1^I24T^ and KRAS^Q61L^ (Fig. 5d, Extended Data Fig. 10a). These values were dramatically higher than for the co-occurrence of any other Q61 mutation with either U2AF1^I24T^ or U2AF1^S34F^. Additionally, there was a strong correlation between the magnitude of exon 3 skipping caused by each Q61 mutation and the likelihood of co-occurrence with U2AF1^I24T^, suggesting that U2AF1^I24T^ mutations are selected for as a means of rescuing normal *KRAS* splicing (Extended Data Fig. 10b).

To further determine whether U2AF1^I24T^ mutations occur secondary to KRAS^Q61^ mutations, we analyzed the VAF of KRAS^Q61R/L^ and U2AF1^I24T^ mutations in 20 pancreatic cancer samples from Foundation Medicine Inc. and AACR Project GENIE which harbored both mutations and had no copy number alterations at the *KRAS* or *U2AF1* loci. Similar to KRAS^G12S^ and U2AF1^S34F^, there was a positive correlation between the two mutations, and the VAF for KRAS^Q61R/L^ was significantly higher than the VAF for U2AF1^I24T^, suggesting that U2AF1^I24T^ occurs as a secondary mutation in KRAS^Q61^-mutant pancreatic cancer (Extended Data Fig. 10c,d).

In total, these results provide a second example of splicing factor mutations that occur as a means of promoting KRAS signaling.

## DISCUSSION

The work presented here describes a mechanism for splicing factor mutation in cancer: certain oncogenic mutations lead to splicing defects that limit their impact on pathogenesis, leading to cascading selection whereby secondary mutations in splicing factors that restore normal splicing are positively selected.

Although U2AF1 mutations have been implicated in several biological pathways that are important in cancer^7,15,18,19^, it has proved difficult to discern the specific selective advantage that these mutations confer in solid tumors. By using a combination of cell modelling and computational investigations of somatic mutations that frequently occurred with mutations in *U2AF1*, we discovered that mutant U2AF1 can regulate the alternative splicing of the *KRAS* oncogene. Our findings suggest that as cancer cells acquire oncogenic KRAS^G12S^ or KRAS^Q61R/L^ mutations, inadvertent exon skipping in *KRAS* leads to reduced KRAS signaling and cell growth. However, the acquisition of U2AF1^S34F^ or U2AF1^I24T^ mutations respectively, reduces exon skipping, driving increased expression of the full-length *KRAS* transcript and increased MAPK signaling and cellular fitness. As a result, *U2AF1* mutations are much more likely to occur in KRAS^G12S^ or KRAS^Q61R/L^-mutant lung or pancreatic adenocarcinomas compared to cancers with other *KRAS* mutations.

Our findings describe a novel mechanism by which mutant U2AF1 may act in cancer: correcting inappropriate transcript splicing that is an unintended and counter-selected consequence of oncogenic mutations. These are the first examples of splicing factor mutations acting to correct splicing errors due to cancer-promoting genomic changes. However, we speculate that this phenomenon may be more widespread. Uncovering additional splicing factor mutations which are selected for as a means of fixing oncogene mis-splicing, if such mutations occur, will require concerted computational and cell modelling efforts.

We observe a clear enrichment of U2AF1^S34F^ and U2AF1^I24T^ mutations in KRAS^G12S^ and KRAS^Q61R/L^-mutant cancers respectively. Additionally, a recent study has found that U2AF1^S34F^ mutations are able to promote cell proliferation in response to overexpression of KRAS^G12V^, perhaps in part by suppressing the expression of inflammatory cytokines^53^. This is consistent with the modest (roughly 2-fold) but statistically significant enrichment of U2AF1^S34F^ mutations in KRAS^G12V^-mutant cancers. However, the majority of *KRAS*-mutant cancers retain wildtype *U2AF1*, and vice versa. This indicates that additional unknown processes drive the positive selection of *U2AF1* mutations in cancer, separate from the ability of mutant U2AF1 to rescue splicing defects in *KRAS*. We and others have shown that U2AF1^S34F^ mutations result in additional alternative splicing and gene expression changes that may affect the growth of cancer cells. Whether these changes act individually or in concert through coordinated gene expression programs remains unknown. Therefore, further analysis of RNA-sequencing datasets, including the one produced here, to uncover unique splicing and gene expression changes induced by specific *U2AF1* variants will be valuable.

Recent work from Kobayashi *et al.* found that KRAS^Q61K^ mutations lead to almost complete *KRAS* exon 3 skipping and early protein termination, resulting in 100% of KRAS^Q61K^-mutant cancers acquiring silent mutations in codon A59 or G60 which restore *KRAS* exon 3 inclusion^49^. We found limited *KRAS* exon 3 skipping in KRAS^Q61K^-mutant TCGA samples, which also show secondary silent mutations^49^. However, KRAS^Q61R^, KRAS^Q61L^ or KRAS^Q61H^ mutations led to varying degrees of *KRAS* exon 3 skipping. Our findings suggest that *KRAS* exon 3 skipping can be tolerated to some degree, but rescue of exon 3 inclusion by means of secondary mutations in *KRAS* or *U2AF1* are strongly selected for depending on the magnitude of exon 3 skipping. As such, KRAS^Q61K^ leads to complete exon 3 skipping and always leads to acquisition of a simultaneous silent mutations^49^. In contrast, other KRAS^Q61^ mutations result in less exon 3 skipping, allowing these mutations to be better tolerated by cancer cells before acquiring secondary U2AF1^I24T^ mutations to drive KRAS signaling.

Our understanding of why certain amino acid substitutions in KRAS are more favored than others is quite limited. Recent work by Huynh and colleagues found that Q61H mutations were over-represented in PDAC, despite Q61R, Q61L and Q61H-mutant cell lines exhibiting similar levels of anchorage-independent growth^54^. We believe that our results partly explain the overabundance of KRAS^Q61H^ mutations, as KRAS^Q61H^ leads to the least amount of exon 3 skipping among the 3 major Q61 mutations. This concept that mutations that result in deleterious exon skipping are less favorable in the absence of secondary mutations may also explain the relative infrequency of KRAS^G12S^ mutations compared to other KRAS mutations at G12.

Our study also provides evidence for another level of regulation for KRAS signaling. Previous studies have reported that KRAS is a relatively weak and poorly optimized oncogene, with a high percentage of rare codons, and weak 3’ splice sites, particularly for exon 2 ^34,43,55^. Our research supports this narrative as exon 2 is readily skipped in the presence of G12S mutations, while exon 3 is also frequently skipped due to multiple mutations at Q61. One possibility for this poor optimization may be that this provides an additional mechanism of regulating KRAS in order to limit oncogene-induced senescence, keeping signaling in a perfect range to induce oncogenesis^55^.

While a ∼15% variation in *KRAS* exon 2 inclusion associated with G12S mutation may seem small, this change was associated with a ∼37% difference in p-Erk levels. Over the course of a developing tumor, this could be sufficient to make significant differences in tumor growth and progression. Additionally, even though a modest decrease in exon 2 inclusion may attenuate the impact of KRAS^G12S^ mutation on cancer cells, we hypothesize that the KRAS mutations that cause exon skipping are nevertheless subject to positive selection as they lock a significant fraction of the protein in a constitutively active state. While genetically engineered mouse models for KRAS^G12S^ are currently unavailable, their development would allow scientists to test these hypotheses and study the impact of combined KRAS and U2AF1 mutations on tumor growth and progression *in vivo*.

Our study also raises the possibility of additional avenues to target KRAS clinically by modulating its splicing. Previous work from the laboratory of Brage Andresen^34,43^ as well as our findings here demonstrate that targeting exon 2 or exon 3 of *KRAS* via SSOs can limit cell growth significantly. Furthermore, we have observed that pharmacological inhibition of the splicing factor SF3B1 also drives *KRAS* exon 2 skipping and suppresses MAPK signaling. As a result, further studies are warranted to test the effectiveness of these therapies in cancers with splice-disrupting KRAS mutations, such as G12S, Q61R and Q61L, as well as in combination with targeted KRAS inhibitors.

Finally, our work provides evidence for a dynamic cancer genome, and offers a new view of evolutionary dependencies where oncogenic mutations are co-selected to drive tumorigenesis^56^. Previous research on *U2AF1* mutations has suggested these occur early in tumor evolution^15,45,46^. However, the variant allele frequencies of KRAS^G12S^ or KRAS^Q61R/L^ are significantly higher than U2AF1^S34F^ or U2AF1^I24T^ in tumors harboring both mutations. Sequential clinical sequencing data are required to provide direct evidence for the secondary nature of U2AF1 mutations. However, our findings are consistent with a model in which a first round of selection events occurs during tumor initiation (KRAS^G12S^ or KRAS^Q61R/L^), that in turn leads to the acquisition of secondary mutational events (U2AF1^S34F^ or U2AF1^I24T^) to compensate for deleterious splicing effects of cancer-causing mutations. Most broadly, our results suggest that first-order cancer-causing mutations may often lead to a cascade of subsequent events whose selection is a response to the first mutation, a process that we term “cascading selection”.

## Methods

### Cell Lines

All cell lines including A549, NCI-H441, NCI-H23, NCI-H2023, Panc 02.13 and 293T cells were obtained from American Type Culture Collection (ATCC). A549, NCI-H441, NCI-H23, and NCI-H2023 cells were cultured in RPMI 1640 medium (Gibco, 11875093) supplemented with 10% fetal bovine serum (Sigma Aldrich, F2442) and 50 μg/ml gentamicin (Gibco, 15750060). Panc 02.13 were cultured in RPMI 1640 medium (Gibco, 11875093) supplemented with 15% FBS, 0.3 mg/ml human recombinant insulin (Sigma Aldrich, 91077C) and 50 μg/ml Gentamicin. 293T cells were cultured in DMEM (Gibco #11965118) supplemented with 10% FBS and 50 μg/ml Gentamicin. Cells were cultured at 37 °C with 5% carbon dioxide. Cells were passaged by washing with phosphate buffered saline (Gibco, 10010023) and trypsinization with 0.25% Trypsin-EDTA (Gibco, 25200114). Cell lines were verified by STR profiling (ATCC, 135-XV) and tested negative for mycoplasma by PCR (ABM, G238).

### Prime Editing

The *U2AF1* and/or *KRAS* alleles of A549 and NCI-H441 cells were modified by twin prime editing using plasmids assembled following the cloning protocol described by Anzalone and colleagues in Supplementary Note 3 ^20^. pU6-tevopreq1-GG-acceptor backbone was a gift from David Liu (Addgene plasmid #174038). Oligonucleotides encoding the spacer sequence, extension template and SpCas9 sgRNA scaffold were ordered from Integrated DNA Technologies (IDT) (Extended Data Table 1) and resuspended in QIAGEN elution buffer at 100 μM concentration. Spacer sequence and extension template oligonucleotides were designed using pegFinder or PRIDICT software^57,58^. Each oligonucleotide pair was annealed by combining 1 μl of top (forward) oligonucleotide, 1 μl of bottom (reverse) oligonucleotide, and 23 μl of annealing buffer (water supplemented with 10 mM Tris-HCl pH 8.5 and 50 mM NaCl), heating at 95 °C for 3 minutes then cooling to 22 °C at 0.1 °C per second. Spacer, extension template and sgRNA scaffold annealed oligonucleotide pairs were then cloned into the GG acceptor backbone via golden gate assembly as described by Anzalone and colleagues in Supplementary Note 3 ^20^.

Cloned plasmids were propagated in NEB Stable Competent *E. coli* (New England BioLabs, C3040) and grown on 50 μg/ml carbenicillin plates (Corning, MT46100RG). Plasmids were extracted using the QIAprep Spin Miniprep Kit (QIAGEN, 27106) and plasmid concentrations were quantified using a Nanodrop ND-1000 Spectrophotometer (Thermo Scientific). Plasmids were then submitted to Quintara Biosciences for Sanger sequencing, or Plasmidsaurus Inc. for whole plasmid long-read sequencing.

Plasmids:

pU6-tevopreq1-U2AF1-S34S-Forward

pU6-tevopreq1-U2AF1-S34S-Reverse

pU6-tevopreq1-U2AF1-S34F-Forward

pU6-tevopreq1-U2AF1-S34F-Reverse

pU6-tevopreq1-U2AF1-S34F(TTC)-Forward

pU6-tevopreq1-U2AF1-S34F(TTC)-Reverse

pU6-tevopreq1-U2AF1-S34Y-Forward

pU6-tevopreq1-U2AF1-S34Y-Reverse

pU6-tevopreq1-U2AF1-S34C-Forward

pU6-tevopreq1-U2AF1-S34C-Reverse

pU6-tevopreq1-U2AF1-S34A-Forward

pU6-tevopreq1-U2AF1-S34A-Reverse

pU6-tevopreq1-U2AF1-S34F-to-WT-Forward

pU6-tevopreq1-U2AF1-S34F-to-WT-Reverse

pU6-tevopreq1-U2AF1-S34F-to-S-Forward

pU6-tevopreq1-U2AF1-S34F-to-S-Reverse

pU6-tevopreq1-U2AF1-S34F-to-F(TTC)-Forward

pU6-tevopreq1-U2AF1-S34F-to-F(TTC)-Reverse

pU6-tevopreq1-U2AF1-S34F-to-Y-Forward

pU6-tevopreq1-U2AF1-S34F-to-Y-Reverse

pU6-tevopreq1-U2AF1-S34F-to-C-Forward

pU6-tevopreq1-U2AF1-S34F-to-C-Reverse

pU6-tevopreq1-U2AF1-S34F-to-A-Forward

pU6-tevopreq1-U2AF1-S34F-to-A-Reverse

pU6-tevopreq1-KRAS-G12S(TCT)-Forward

pU6-tevopreq1-KRAS-G12S(TCT)-Reverse

Twin prime editing (TwinPE) was carried out by transfecting human cell lines with pCMV-PEMax-P2A-BSD (Addgene plasmid #174821) alongside 2 complementary epegRNAs cloned into the pU6-tevopreq1-GG-acceptor backbone (Addgene plasmid #174038)^21^. Cells were plated at 2 × 10^5^ cells per well in a 6-well plate, and 24 hours after plating transfections were conducted using the Mirus Bio TransIT-X2 Dynamic Delivery System (Mirus Bio LLC, MIR6004) according to the manufacturer’s recommended protocol for 6-well plates. A549 cells were treated with 5 μl of TransIT-X2 reagent in 6 well plates, while NCI-H441 cells were treated with 7.5 μl of TransIT-X2 reagent. 500 ng each of the forward and reverse epegRNA vectors were used, along with 1.5 μg of pCMV-PEMax-P2A-BSD. Cells were then treated with 10 mg/ml Blasticidin S HCl (Gibco, A1113903) 24-hours post-transfection and changed to normal media 72 hours post-transfection.

### Single Cell Cloning

Cells were counted 1 week after prime editing, and 10,000 cells were plated in well A1 of a 96-well plate before being subjected to array dilution^59^. After 2-4 weeks, wells with single colonies were trypsinized and half of the cell volume was passaged in 24-well plates. 1 mL PBS was added to the remaining half of cells before being centrifuged for 5 minutes at 340 × *g*. The PBS was aspirated, and the cells were resuspended in 10 μl Elution Buffer (QIAGEN) and boiled at 100°C for 15 minutes. Samples were then placed on ice for 3 minutes before being pelleted by centrifugation at 20,000 × *g*. for 1 minute. PCR amplification of the desired locus was performed with 12.5 μl Q5 High-Fidelity 2X Master Mix (New England BioLabs, M0494S) along with 7.5 μl of the isolated supernatant as template, and 2.5 μl each of 5 μM forward and 5 μM reverse PCR primers (Extended Data Table 5). Samples were then submitted to Quintara Biosciences for Sanger sequencing.

### RNA-Sequencing

RNA for each cell clone was extracted using the RNeasy Plus Mini Kit (QIAGEN, 74134) as per the manufacturer’s instructions. RNA concentration was quantified using the Qubit RNA Broad Range Quantification Assay Kit (Invitrogen, Q10210) according to the manufacturer’s protocol. RNA was then submitted to Novogene for mRNA-Sequencing including the following steps described in this paragraph. Sample quality was confirmed to have a RIN score >9 by BioAnalyzer. Messenger RNA was purified from total RNA using poly-T oligo-attached magnetic beads. After fragmentation, the first strand cDNA was synthesized using random hexamer primers, followed by the second strand cDNA synthesis using dTTP. The samples were then processed by end repair, A-tailing, adapter ligation, size selection, amplification, and purification. The library quality was checked with Qubit and real-time PCR for quantification and BioAnalyzer for size distribution detection. Paired-end clean reads were aligned to the reference genome GRCh38^60^ using Hisat2 (v2.0.5)^61^. FeatureCounts (v1.5.0-p3) was used to count the reads numbers mapped to each gene^62^.

Alternative splicing events were identified using rMATS (4.1.0)^63^ and defined as significant according to an FDR < 0.05. Sashimi plots were viewed and generated using Integrative Genomics Viewer (v2.16.10)^64^. Filtering for known oncogenes and tumor suppressors was performed by examining genes that were defined as cancer genes by at least 4 sources in OncoKB^27,28^.

*KRAS* exon 2 read coverage and fraction of mutant reads for *U2AF1* were analyzed using Integrative Genomics Viewer (v2.16.10)^64^. The overlap of significant alternative splicing events between U2AF1^S34F^-mutant A549 cells and human lung adenocarcinomas was quantified by performing a Fisher’s exact text on alternative splicing events (p<0.05 by rank-sum test) present or absence in each dataset as determined by rMATS (A549 cells) or SplAdder (TCGA)^24^.

To identify mutant U2AF1 3’ splice site sequence preferences, significant exon splicing events were first determined using rMATS. The regions of interest were set as 10 bp preceding and following the first base of each skipped exon, as well as 10 bp preceding and following the first base of each included exon. The coordinates of these genomic regions were recorded in BED file format. Sequences corresponding to these regions were extracted from the Hg38 human reference genome fasta file using the getfasta function from bedtools (v2.26.0)^65^, and graphics were generated using Weblogo (v.3.7.11)^66^.

For long-read RNA-Sequencing, RNA was isolated as above and submitted to Broad Clinical Labs (Broad Institute). An aliquot of 300ng of total RNA was used as the input for Kinnex™ full length cDNA synthesis and amplification (PacBio Iso-Seq express 2.0 kit, 103-071-500). The amplified cDNA was then quantified by Qubit and Tapestation (Qubit dsDNA HiSens, QUBDSDNA500KT, and HiSense D5000 ScreenTape, 50675592) and barcoded cDNA was then pooled before proceeding to 8-fold Kinnex PCR (PacBio Kinnex PCR 8-fold kit, 103-071-600). Samples then were programmatically concatenated into single molecules optimal for long-read sequencing (Kinnex full-length RNA kit, 103-072-000). Success of array formation was evaluated through qubit quantification and Agilent TapeStation fragment size QC (Qubit dsDNA HiSens, QUBDSDNA500KT, and GenomicDNAScreenTape, 50675366). Each pooled Kinnex library underwent sequencing preparation using the Revio polymerase kit (Pacific Biosciences, 102-739-100). Using the Sample Setup page in SMRTLink, the appropriate volumes of each reagent was calculated by factoring in the insert size, sample concentration, and target plate loading concentration of 150pM. The run design was set up with the Application Type “Kinnex full-length RNA” and Library Type “Kinnex.” Samples were loaded onto the sequencer within 24 hours of completed sequencing preparation and were sequenced using the 24-hour movie setting.

Following sequencing, unaligned BAM files were first converted to FASTQ format with SAMtools fastq^67^. Samples were then aligned to the primary GRCh38.p14^60^ assembly with Minimap2^68^ using the “-ax splice:hq -uf” parameters. The alignments were then sorted and indexed with samtools index and samtools sort, respectively, to create aligned BAM files. For transcript quantification and discovery, Isoquant^69^ was run on the aligned BAM files individually with transcript discovery enabled using the Gencode.v48 primary annotation^70^. Transcript discovery was performed on each sample individually, rather than through a joint transcript model of all samples or a subset of samples, to provide us with high sensitivity to novel transcripts. Stringtie –merge^71^ was then used to extend the Gencode.v48.primary annotation with the unique set of novel transcripts that were identified. Two novel KRAS transcripts (MSTRG14141.1 and MSTRG14141.13) were identified, and then Isoquant was re-run without transcript discovery to quantify all samples using the merged annotation.

### RT-PCR quantification of *KRAS* exon 2 or exon 3 skipping

RT-PCR primers to detect *KRAS* exon 2 or exon 3 skipping were designed using Primer-BLAST^72^, with the forward and reverse primers being present on exons flanking the exon of interest (Extended Data Table 5). Cells were trypsinized and centrifuged at 340 × *g* for 5 min, followed by RNA extraction using RNeasy Plus Mini Kit (QIAGEN, 74134) as per manufacturer’s protocol. RNA concentrations were quantified with the Qubit RNA BR Assay Kit (Invitrogen, Q10210), and 100 ng of RNA was used to make cDNA using the SuperScript IV VILO Master Mix (Invitrogen, 11756050). The samples were PCR amplified using Phusion Plus PCR Master Mix (Thermo Scientific, F631S), using a 60°C annealing temperature per the manufacturer’s 3-step protocol. The PCR products were run on a 2% agarose gel and imaged with a ChemiDoc MP (Bio-Rad Laboratories, 12003154). Band intensities were quantified using the Analyze Gels feature in Fiji (ImageJ). Briefly, horizontal boxes were drawn around the top (exon included) and bottom (exon skipped) bands, and the intensity of each band was quantified. The sum of the band intensities was then calculated, and each band intensity was divided by the total intensity to give the fraction of exon inclusion.

### qRT-PCR quantification of *KRAS* exon 2 skipping

qRT-PCR primers to detect *KRAS* exon 2 skipping and inclusion were designed using Primer-BLAST^72^, with the forward primer spanning either the exon 1-exon 3 junction (for skipping) or the exon 1-exon 2 junction (for inclusion) (Extended Data Table 5). qRT-PCR primers to detect Beta-Actin expression for an internal control were also designed using Primer-BLAST^72^ (Extended Data Table 5). Cells were trypsinized and centrifuged at 340 × *g* for 5 min, followed by RNA extraction using the RNeasy Plus Mini Kit (QIAGEN, 74134) as per the manufacturer’s instructions. RNA concentrations were quantified with the Qubit RNA BR Assay Kit (Invitrogen, Q10210), and 100 ng of RNA was used to make cDNA using the SuperScript IV VILO Master Mix (Invitrogen, 11756050). cDNA was diluted 1:25 and 1.5 μl was used as input along with 0.5 μl each of forward and reverse primers at a concentration of 100 nM each, as well as 2.5 μl of Power SYBR Green PCR Master Mix (2X) (Applied Biosciences, 4368577) for qRT-PCR following the manufacturer’s two-step protocol. The fold change of *KRAS* exon 2 skipping or inclusion was quantified using the 2^-ΔΔCt^ method by first normalizing expression to Beta-actin, then normalizing all samples to U2AF1^S34S^ sample 1, before quantifying the fold change.

### Modeling U2AF1 binding to *KRAS* RNA

To model U2AF1 binding to *KRAS* RNA, RBPSuite (v1.0)^37^ was used to generate predicted U2AF1 binding scores. Human was used as the species, linear RNA was selected as RNA type, and U2AF1 was selected as the specific model. A 46 bp RNA sequence centered at the 3’ splice site of *KRAS* exon 2, or a 34 bp RNA sequence centered at wildtype *KRAS* amino acid G12, or mutant *KRAS* G12S was used as the input sequence.

Exon 2 3’ splice site RNA sequence input: CATTTTCATTATTTTTATTATAAGGCCTGCTGAAAATGACTGAATA

Wildtype G12 RNA sequence input: GTGGTAGTTGGAGCTGGTGGCGTAGGCAAGAGTG

G12S RNA sequence input: GTGGTAGTTGGAGCTAGTGGCGTAGGCAAGAGTG

### Modeling steric hindrance and *KRAS* exon 2 skipping using dCasRx

pXR002: EF1a-dCasRx-2A-EGFP and pXR003: CasRx gRNA cloning backbone were gifts from Patrick Hsu (Addgene plasmid # 109050 and 109053)^40^. dCasRx sgRNAs were designed using the cas13design tool (https://cas13design.nygenome.org)^73,74^, ordered from IDT and resuspended in QIAGEN elution buffer (Extended Data Table 3). Each oligonucleotide pair was annealed by combining 1 μl of top (forward) oligonucleotide, 1 μl of bottom (reverse) oligonucleotide, and 23 μl of annealing buffer (water supplemented with 10 mM Tris-HCl pH 8.5 and 50 mM NaCl), heating at 95 °C for 3 minutes then cooling to 22 °C at 0.1 °C per second.

Annealed oligonucleotide pairs were then cloned into the CasRX gRNA cloning backbone via golden gate assembly, using BbsI-HF (NEB R3539S)^20^. Cloned plasmids were propagated in NEB Stable Competent *E. coli* (New England BioLabs, C3040) and grown on 50 μg/ml carbenicillin plates (Corning, MT46100RG). Plasmids were extracted using the QIAprep Spin Miniprep Kit (QIAGEN, 27106) and plasmid concentrations were quantified using a Nanodrop ND-1000 Spectrophotometer (Thermo Scientific). Plasmids were then submitted to Quintara Biosciences for Sanger sequencing, or Plasmidsaurus Inc. for whole plasmid long-read sequencing.

Plasmids:

pXR003-KRAS-Exon2-3’SS-1

pXR003-KRAS-Exon2-3’SS-2

pXR003-KRAS-G12-1

pXR003-KRAS-G12-2

pXR003-KRAS-Downstream-4

pXR003-KRAS-Downstream-5

To test the impact of dCasRx binding to different locations on the *KRAS* exon 2 RNA, NCI-H2023 with wildtype *KRAS* were plated at 200,000 cells per well on 6-well plates. Cells were transfected 24 hours later with 5 μg EF1a-dCasRx-2A-EGFP and 5 μg dCasRx sgRNA in 500 μl Opti-MEM with 20 μl TransIT-X2. 48 hours post-transfection, cells were trypsinized and centrifuged at 340 × *g* for 5 min, followed by RNA extraction using RNeasy Plus Mini Kit (QIAGEN, 74134). 100 ng of RNA was used to make cDNA using the SuperScript IV VILO Master Mix (Invitrogen, 11756050). Exon 2 skipping was then quantified by RT-PCR, and the PCRs were run on a 2% agarose gel.

### Splicing Reporter Minigene

The pSpliceExpress system used for splicing reporter minigene experiments was a gift from Stefan Stamm (Addgene plasmid #32485)^41^. Wildtype *KRAS* Exon 2 and Exon 3 DNA (containing 500 bp of intronic sequence upstream and downstream of each exon) was synthesized as gBlocks by Integrated DNA Technologies (IDT) and then cloned into the pSpliceExpress via Gibson assembly. Simply, the gBlock was amplified using 2X Phusion Plus PCR Master Mix (Thermo Fisher, F631S) and primers with overlapping sequences to the pSpliceExpress backbone to insert the fragment between rat insulin exons 2 and 3, following the manufacturer’s recommended 3-step protocol. The pSpliceExpress vector was also linearized by PCR using 2X Phusion Plus PCR Master Mix (Thermo Fisher, F631S) following the manufacturer’s recommended 3-step protocol. Gibson assembly primers were designed using the NEBuilder Assembly Tool (NEB). 10% of each PCR product was run on a 1% agarose gel to confirm the correct size, and the remaining product was PCR purified using the QIAquick PCR & Gel Cleanup Kit (QIAGEN, 28506). Gibson assembly was then carried out using PCR products for pSpliceExpress and *KRAS* exon 2 or exon 3, and NEBuilder Hifi DNA Assembly master mix (NEB, E2621S) using the manufacturer’s recommended protocol. The cloned plasmids were propagated in NEB Stable Competent E. coli (New England BioLabs, C3040) and grown on 50 µg/ml carbenicillin plates (Corning, MT46100RG). Plasmid extraction was completed with the QIAprep Spin Miniprep Kit (QIAGEN, 27106) and plasmid concentrations were measured using a Nanodrop ND-1000 Spectrophotometer (Thermo Scientific) and submitted to Quintara Biosciences for whole plasmid sequencing.

*KRAS* exon 2 and exon 3 mutations were introduced using a modified site-directed mutagenesis protocol^75^. Site directed mutagenesis primers were designed using PrimerX (https://www.bioinformatics.org/primerx/) and site-directed mutagenesis was performed by PCR amplification with Platinum SuperFi II Master Mix (Thermo Fisher, 12368010) following the manufacturer’s protocol. The reaction was then treated with DpnI (NEB, R0176S) for 1 hour at 37°C before bacterial transformation.

To study levels of KRAS exon 2 and exon 3 skipping cells, 293T cells were seeded at 100,000 cells per well in a 24-well plate. After 24 hours, cells were transfected with 1 μg pSpliceExpress plasmid in 100 uL of Opti-MEM I (Gibco, 31985062) with 3 µl of Mirus Bio TransIT-X2 Dynamic Delivery System (Mirus Bio LLC, MIR6004). 24 hours after transfection, cells were trypsinized and centrifuged at 340 x g for 5 min, followed by RNA extraction using RNeasy Plus Mini Kit (QIAGEN, 74134). 200 ng of RNA was used for cDNA production using the SuperScript IV VILO Master Mix (Invitrogen, 11756050). Exon skipping was then quantified by RT-PCR using Phusion Plus PCR Master Mix (ThermoFisher, F632S) and primers specific to the pSpliceExpress system^76^ (Extended Data Table 5). PCRs were run on a 2% agarose gel and band intensities were quantified using the Analyze Gels feature in Fiji (ImageJ) as described above.

Plasmids:

pSplice-Express-KRAS-Exon2

pSplice-Express-KRAS-G12S

pSplice-Express-KRAS-G12S(TCT)

pSplice-Express-KRAS-G12A

pSplice-Express-KRAS-G12C

pSplice-Express-KRAS-G12D

pSplice-Express-KRAS-G12V

pSplice-Express-KRAS-G10R(AGA)

pSplice-Express-KRAS-G10R(CGA)

pSplice-Express-KRAS-A11A(GCA)

pSplice-Express-KRAS-A11A(GCC)

pSplice-Express-KRAS-G13S(AGC)

pSplice-Express-KRAS-G13S(TCC)

pSplice-Express-KRAS-F28L(TTA)

pSplice-Express-KRAS-F28L(TTG)

pSplice-Express-KRAS-Exon3

pSplice-Express-KRAS-Q61R

pSplice-Express-KRAS-Q61L

pSplice-Express-KRAS-Q61H

pSplice-Express-KRAS-Q61P

pSplice-Express-KRAS-Q61K

pSplice-Express-KRAS-GQ60GK

### Flow Cytometry

For flow cytometry experiments to measure p-Erk levels, 500,000 cells were first plated in 6 well plates. 24 hours after plating, cells were deprived of serum and then were collected 16 hours later by trypsinization using 500 μl of trypsin. An equal volume of pre-warmed BD Phosflow Fix Buffer I (BD Biosciences, 557870) was added directly to the plate, cells were incubated at 37C for 10 minutes before being collected and centrifuged at 340 × *g* for 5 min. Cells were washed with PBS and permeabilized with 1 ml ice cold BD Phosflow Perm Buffer III (BD Biosciences, 558050) and incubated on ice for 30 minutes. Cells were counted, and equal numbers of cells were washed twice with staining buffer (1X PBS, 1% FBS and 0.09% sodium azide) before being resuspended in 50 μl of staining buffer with BD Phosflow PE-Cy7 anti-Erk1/2 (pT202/pY204) antibody (BD Biosciences, 560116) at a 1:5 dilution. Cells were stained for 30 minutes at room temperature in the dark, then washed with PBS and resuspended in PBS at a final concentration of 1 million cells per ml. Samples were then analyzed on a BD LSRFortessa Cell Analyzer (BD Biosciences) at the Dana-Farber Cancer Institute Flow Cytometry Core.

### Pladienolide B Treatment

To determine the impact of SF3B1 inhibition on *KRAS* exon 2 skipping and MAPK signaling, 500,000 NCI-H2023 cells were first plated in 6 well plates. 24 hours later, cells were treated with DMSO or Pladienolide B at concentrations of 100 nM, 50 nM, 10 nM, 5 nM or 1 nM, being sure that all samples received equal total amounts of DMSO to control for potential DMSO-mediated effects. For flow cytometry measurement of p-Erk, cells were also changed to serum free media. 20 hours after Pladienolide B treatment, cells were collected for RNA isolation or flow cytometry. *KRAS* exon 2 skipping and p-Erk levels were measured by RT-PCR and flow cytometry respectively, as described above.

### Splice Switching Oligonucleotide Treatment

For *KRAS* exon 2 skipping, SSOs designed and validated by the laboratory of Brage Andresen were used^43^. For *KRAS* exon 3 skipping, SSOs were designed using the eSkip-Finder application^77^, using 2’OMe chemistry and a length of 25 bp. SSO sequences are available in Extended Data Table 4. NCI-H23 cells were plated at 2 × 10^6^ cells per 10 cm plate. 24 hours after plating, cells were transfected using Transit-X2 transfection reagent according to the manufacturer’s recommended protocol for 10 cm plates, with 45 μl of TransIT-X2 reagent. Each well was treated with 120 nM total SSO, comprising of a range of concentrations of either scramble SSO control, or *KRAS* exon 2 or exon 3 skipping SSO (120 nM scramble SSO alone, 6 nM *KRAS* SSO with 114 nM scramble SSO, 12 nM *KRAS* SSO with 108 nM scramble SSO, 30 nM *KRAS* SSO with 90 nM scramble SSO, 60 nM *KRAS* SSO with 60 nM scramble SSO, or 120 nM *KRAS* SSO alone) to keep the total amount of SSO delivery the same between conditions^34,43^. Cells were plated for cell growth assays 24 hours after transfection, while cells were collected for RNA or protein 72 hours following transfection.

### Immunoblot Analysis

For immunoblot analyses, cells were deprived of serum for 16 hours before cell collection to reduce the impact of extracellular growth factor stimulation on MAPK signaling. Cells were lysed with RIPA Buffer (Millipore Sigma, R0278-50ML) containing Halt Protease and Phosphatase Inhibitor Cocktail (100X) (Thermo Scientific, 78440) and quantified with the Pierce BCA Protein Assay Kit (Thermo Scientific, 23225). Following addition of NuPAGE 4X LDS Sample Buffer (Invitrogen, NP0008) and NuPAGE 10X Sample Reducing Agent (Invitrogen, NP0009), the samples were run on a NuPAGE 4-12% Bis-Tris, 1.5 MM, Mini Protein Gel (Invitrogen, NP0335BOX) in NuPAGE 20X MOPS SDS Running Buffer (Invitrogen, NP0001) at 125V for 105 min on ice. The gel was transferred on a nitrocellulose membrane in NuPAGE 20X Transfer Buffer (Invitrogen, NP00061) at 70V for 2 hours on ice. Blocking and Immunostaining for HSP90 (BD Biosciences, 610418, 1:10,000), p-Erk (Cell Signaling, 4370S, 1:2000), Total-Erk (Cell Signaling, 9102S, 1:2000), p-Akt (Cell Signaling, 4060S, 1:1000), Total-Akt (Cell Signaling, 58295S, 1:500) and KRAS (Invitrogen, 11H35L14, 1:2500) were done in Intercept (TBS) Blocking Buffer (LI-COR Biosciences, 927-60001), with 0.1% Tween-20 for primary and secondary antibody blockings. LI-COR Odyssey Classic Imaging System (LI-COR Biosciences, ODY-9120) was used to image the blots.

### Cell Proliferation Assays

Cells were plated on ultra-low attachment plates (Corning, 3474) at 1,000 cells per well in 100 μl of media for each desired time point to measure growth. At each time point, cells were moved to white-walled 96-well plates (Corning, 3903). 100 μl of CellTiter-Glo Luminescent Cell Viability Assay (Promega, G7572) diluted 1:1 in PBS was then added to each well. After a 10-minute incubation at room temperature while rocking, the plates were analyzed using a SpectraMax M5 (Molecular Devices) plate reader using the included CellTiter-Glo protocol with a luminescence integration time of 500 ms. Luminescence values were normalized to data points from the earliest time point.

### Cloning of lentiviral vectors

U2AF1^S34F^ cDNA was synthesized by IDT and subsequently cloned into the pLX301 backbone (Addgene #25895) via Gibson assembly. In brief, the U2AF1^S34F^ cDNA was amplified using 2X Phusion Plus PCR Master Mix (Thermo Fisher, F631S) and primers with overlapping sequences to the pLX301 backbone following the CMV promoter. Gibson assembly primers were designed using the NEBuilder Assembly Tool (NEB) and contained a FLAG tag sequence added to the C-terminal end of U2AF1. The PCR product was run on a 1% agarose gel and gel purified using the QIAquick PCR & Gel Cleanup Kit (QIAGEN, 28506). The pLX301 backbone was linearized via restriction digested with BsrGI-HF (NEB, R3575S) for 1 hour at 37°C and then run on a 1% agarose gel and gel purified. Gibson assembly was then carried out using linearized pLX301, U2AF1 PCR product, and NEBuilder Hifi DNA Assembly master mix (NEB, E2621S) using the manufacturer’s recommended protocol.

The vector was then mutated to express wildtype U2AF1 and U2AF1^I24T^ via sequential site-directed mutagenesis. First, pLX301-U2AF1^S34F^-FLAG was mutated to pLX301-U2AF1-FLAG, before being mutated again to form pLX301-U2AF1^I24T^-FLAG. Site directed mutagenesis primers were designed using PrimerX (https://www.bioinformatics.org/primerx/) and site-directed mutagenesis was performed by PCR amplification with Platinum SuperFi II Master Mix (Thermo Fisher, 12368010) followed by restriction digestion with DpnI (NEB, R0176S) for 1 hour at 37°C before bacterial transformation.

Plasmids:

pLX301-U2AF1-WT-FLAG

pLX301-U2AF1-S34F-FLAG

pLX301-U2AF1-I24T-FLAG

### Lentivirus Production and Transduction

Lentivirus was produced as described previously^78^. Briefly, 8 × 10^6^ 293T cells were plated on 15 cm plates coated with 0.1% gelatin. 24 hours after plating, cells were transfected with 10 μg pLX301 lentiviral vector, 7.5 μg delta 8.9 plasmid, and 2.5 μg pCMV-VSV-G along with 80 μl PEIMax (Polysciences, 24765) in 1 ml of Opti-MEM I (Gibco, 31985062). 24 hours after transfection, the media was replaced with fresh DMEM supplemented with 25 mM HEPES (Gibco, 15630-080) and 3 mM caffeine (Sigma, C0750). Lentivirus-containing supernatant was collected from the cells at 48 and 72 hours following transfection and filtered through 0.45-μm filters (Thermo Scientific 723-2545).

Lentivirus was then concentrated by centrifugation through a sucrose cushion^79^. A 10% sucrose solution (50 mM Tris-HCl,100 mM NaCl, 0.5 mM EDTA) was carefully pipetted underneath the lentivirus-containing media at a 4:1 ratio of lentivirus-containing media to sucrose solution. The lentivirus was then centrifuged at 10,000 × g for 4 hours at 4°C. Media was aspirated and the pellet was resuspended in 100 μl of PBS.

For Panc 02.13 cell transduction, 35,000 cells were plated in 300 μl insulin-containing RPMI in 48-well plates. 24 hours after plating, cells were transduced via spinfection. Briefly, fresh RPMI containing 8 μg/ml polybrene (Santa Cruz Biotechnology, sc-134220) was added to each well. The titer of each lentivirus was estimated using Lenti-X GoStix Plus (Takara, 631280), and equal amounts of lentivirus was delivered to each well. The 48-well plate was then centrifuged at 1,000 × g for 2 hours at 30°C. 24 hours after spinfection, fresh RPMI containing 2 μg/ml puromcycin (Gibco, A1113803) was added to each well. 48 hours after puromycin treatment, cells were trypsinized and collected for RNA isolation and RT-PCR.

### Genomic Data

All data from the AACR Project GENIE Consortium^11^ were obtained through the dedicated cBioPortal website^29^. Additional lung adenocarcinoma and pancreatic cancer sequencing data were provided by Foundation Medicine Inc., comprising 62,009 lung cancer cases and 31,530 pancreatic cancer cases with tissue biopsy-based comprehensive genomic profiling using FoundationOne^®^/ FoundationOne^®^CDx during routine clinical care^26^. In these cohorts, co-occurrence of *U2AF1* and *KRAS* mutations, as well as a comparison of variant allele frequencies for mutant *U2AF1* and *KRAS* in patients with both alterations were assessed.

Alternative splicing data from CCLE was obtained from Ghandi *et al*.^35^, and *KRAS* and *U2AF1* mutational data was obtained via cBioPortal^29^. Alternative splicing data from TCGA was obtained from Kahles *et al.*^24^, and *KRAS* and *U2AF1* mutation data was obtained via cBioPortal^29^.

### Statistical Analysis

All statistical analyses were performed in GraphPad Prism 10 except where noted. Means and error bars (standard deviation) are plotted for all analyses. Measurements were taken from distinct samples (biological replicates) unless noted. Alternative splicing events were analyzed, and FDR q values were calculated using rMATS^63^ for short-read RNA-sequencing, and rMATS-long^63^ for long-read RNA-Sequencing. A simple linear regression was performed for the correlation between the fraction of exon 2 inclusion and the variant allele frequency for KRAS^G12S^ in TCGA, the correlation between the variant allele frequency of KRAS^G12S^ and U2AF1^S34F^ in the Foundation Medicine Inc. dataset, the correlation between the variant allele frequency of KRAS^Q61R/L^ and U2AF1^I24T^ in the Foundation Medicine Inc. and AACR Project GENIE datasets, and the correlation between the odds ratio of co-occurrence with U2AF1^I24T^ for each KRAS mutation at Q61 and the fraction of *KRAS* exon 3 inclusion for each mutation. A nonlinear (semi-log) fit was performed to analyze the correlation between the percentage of patients with U2AF1^S34F^ mutations and the odds ratio of a smoker developing the disease, and the correlation between the number alternatively spliced genes and the relative frequency of each U2AF1 mutation across all cancers.

To quantify the fraction of mutant reads for each *U2AF1* variant, the fraction of reads with each individual splicing event across cells with different *U2AF1* variants, the fraction of reads with *KRAS* exon skipping and *KRAS4A* isoform usage in CCLE and TCGA data, and the fraction of *KRAS* exon 2 inclusion in CCLE and TCGA samples with and without U2AF1^S34F^ and KRAS^G12S^ mutations, a one-way ANOVA with multiple comparisons using Dunnett correction was performed. The two-sided Student’s t-test was used to analyze the fraction of RNA-sequencing reads showing *KRAS* exon 2 skipping in backedited A549 cells and engineered NCI-H441 cells, the fraction of *KRAS* exon 2 skipping in U2AF1^S34F^-mutant A549 cells by RT-PCR and qRT-PCR, and p-Erk levels in A549 cells with U2AF1^S34S^, U2AF1^S34F^ and KRAS^G12S(TCT)^ mutations. For the comparison of the variant allele frequency of KRAS^G12S^ and U2AF1^S34F^, and KRAS^Q61R/L^ and U2AF1^I24T^, a two-sided paired t-test was used.

A two-sided Fisher Exact test was used to compare the overlap in splicing events between engineered U2AF1^S34F^-mutant A549 cells and U2AF1^S34F^-mutant lung adenocarcinoma samples, for analysis of the frequency of mutations in U2AF1^S34F^-mutant lung adenocarcinoma compared to all other lung adenocarcinoma cases, for the calculation of co-occurrence between U2AF1 and KRAS mutations from Foundation Medicine Inc. and AACR Project GENIE data, and for the identification of additional somatic mutations which co-occur with splicing factor mutations.

## Data Availability

Short- and long-read RNA-sequencing data will be available in the NCBI Gene Expression Omnibus (GEO). Unique plasmids will be available in Addgene. All other data generated and analyzed during this study are included in the manuscript. Requests for further information should be directed to the lead contact.

## Code Availability

This paper does not report original code.

## Acknowledgements

We wish to thank Mitchell Leibowitz, Owen Hirschi, and all the members of the Meyerson laboratory for their advice and technical assistance; Mitchell suggested the steric hindrance experiment and Owen suggested the use of splicing reporter minigenes. Thank you to Leslie Gaffney for assistance with figures. Thank you to Peter Chen and David Liu for their guidance with prime editing approaches and for sharing detailed protocols with us. Thank you to Andrew Aguirre, Ben Lampson and Colleen Harrington for helpful discussions. This research was supported by Damon Runyon Cancer Research Foundation awards (D.M.W. DRG-2449-21 and A.B.D. DRG-2504-23.), an NIH R35 CA197568 grant (M.M.) and the American Cancer Society Research Professorship (M.M.).

## Contributions

D.M.W. and M.M. conceived of the study. D.M.W., K.C. and D.D. designed and performed most of the experiments and analyzed most of the data. S.S. and G.F. performed and analyzed the genomics data from Foundation Medicine Inc. I.T.-H.L., A.B.D., J.M.H., E.S., K.X.J. and A.A.G. performed additional genomic analyses. D.M.W., K.C. and M.M. wrote the manuscript. All authors read and edited the manuscript.

## Ethics Declarations

S.S. and G.F. are employees of Foundation Medicine Inc., a subsidiary of Roche, and have stock ownership in Roche. M.M. consults for and holds equity in Bayer and Delve Bio; holds equity in Isabl and Karyoverse; is an inventor on patents licensed to Bayer and Labcorp; and receives research support from Bayer and Janssen, all outside the scope of the current work. M.M. was also a founder of Foundation Medicine with shares sold to Roche but has no continued financial relationship with the company at the time of manuscript submission.

---

**Extended Data Fig. 1:**
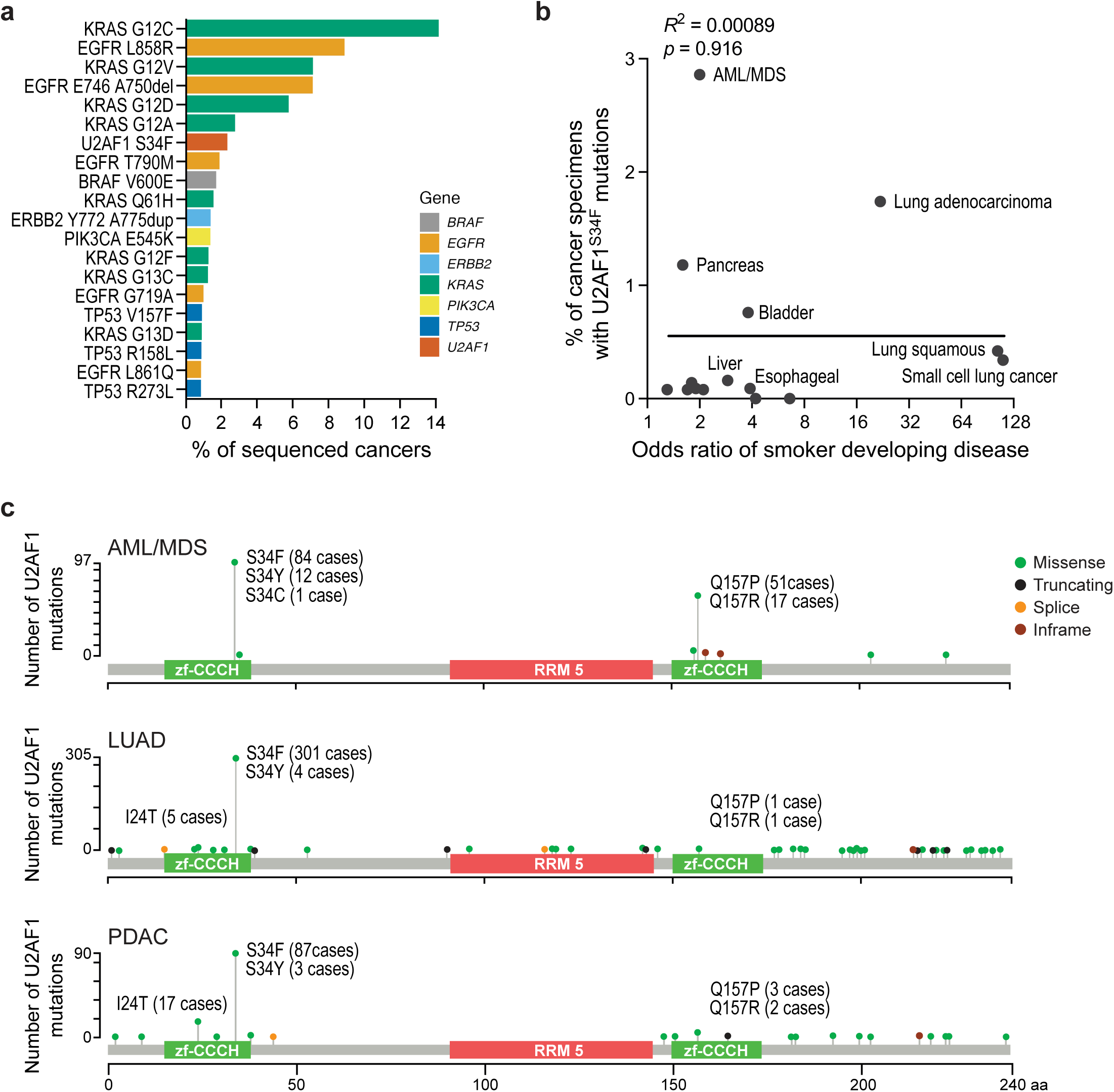
U2AF1^S34F^ mutations are over-represented in human lung adenocarcinomas. a) Top 20 most frequent hotspot mutations in human lung adenocarcinomas based on an analysis of data from AACR Project GENIE (v15.0)^11^. b) Comparison of the percentage of cancer specimens with U2AF1^S34F^ mutations across cancer types (AACR Project GENIE v15.0)^11^ compared to the odds ratio of a smoker developing that disease^14^ (F=0.012, DFn=1, DFd=13,R^2^=0.00089, p=0.92). c) Lollipop plots of mutations identified in the *U2AF1* gene in acute myeloid leukemia/myelodysplastic syndrome (n=181 patients), pancreatic ductal adenocarcinoma (n= 131 patients) and lung adenocarcinoma (n=380 patients) based on data from AACR Project GENIE (v16.0)^11,29^.

**Extended Data Fig. 2:**
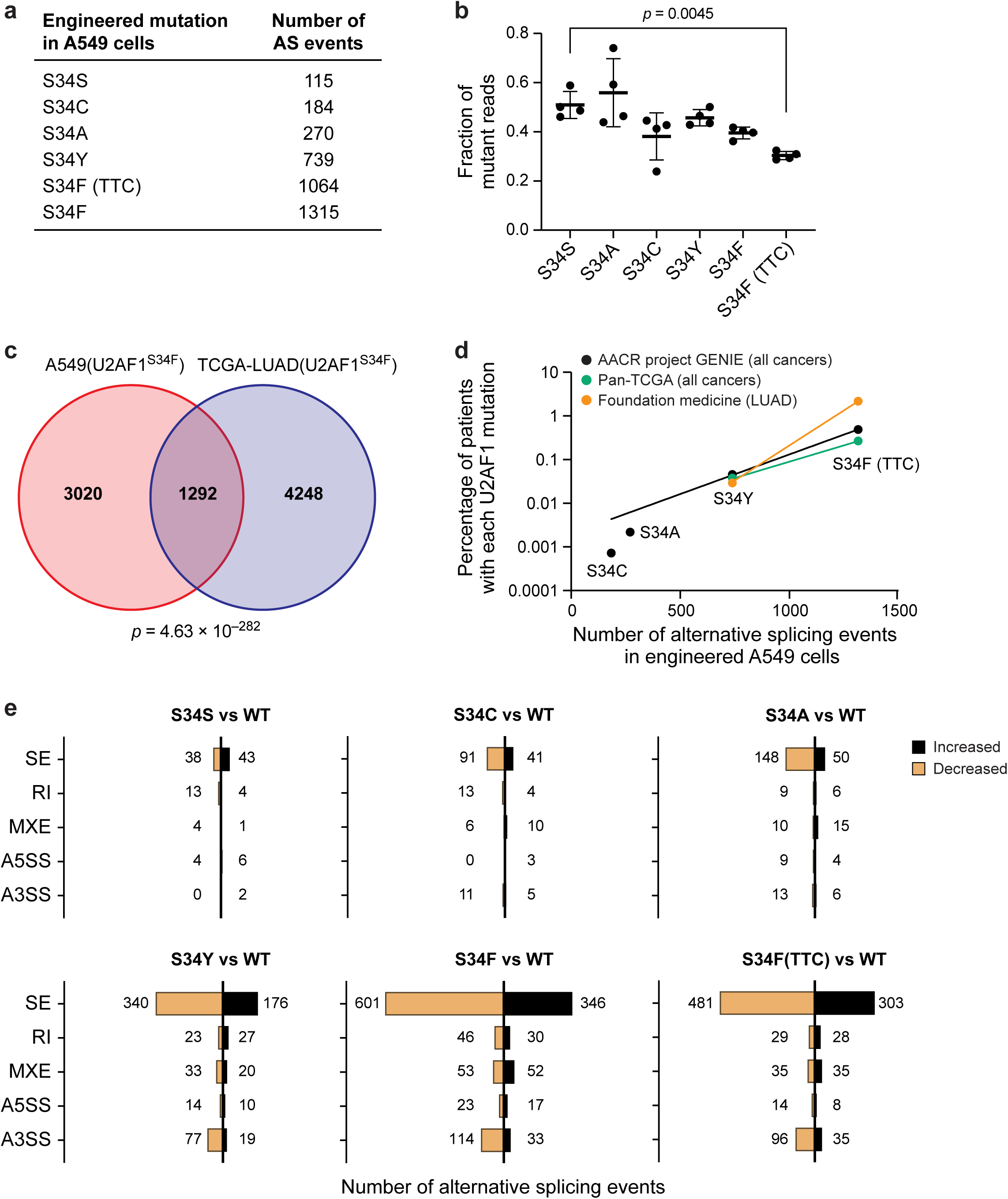
Alternative splicing analysis of A549 cells with varying U2AF1 amino acid substitutions. a) Number of significant alternative splicing events identified in the context of each engineered *U2AF1* mutation in A549 cells (n=4 clones each). Significant alternative splicing events were identified by comparing each variant to transcripts from parental A549 cells using rMATS (v4.1.0) and defined by an FDR q value < 0.05. b) Fraction of RNA-sequencing reads for each engineered *U2AF1* variant sequence relative to wildtype *U2AF1* in A549 cells (n=4 clones each, q ratio=3.90, DF=18, p=0.0045 for S34S vs. S34F(TTC)). c) Overlap of significant alternative splicing events between mRNA from A549 cells with an engineered U2AF1^S34F^ mutation (n=4 clones compared to 4 parental A549 cell controls) and mRNA from U2AF1^S34F^-mutant lung adenocarcinoma patient samples from TCGA^24^ (n=7 cases compared to 100 randomly sampled controls with wildtype *U2AF1*) (odds ratio=4.18, p=4.63x10^-282^). d) Correlation between the number of alternative splicing events that occur in the presence of each amino acid substitution in A549 cells and the frequency of each substitution across all human cancers as determined by the AACR Project GENIE dataset (v15.1), TCGA Pan-Cancer Atlas, or Foundation Medicine Inc. LUAD data^11,25,26^. e) Number of significant alternative splicing events for A549 cells with S34S (n=4 clones), S34C (n=4 clones), S34A (n=4 clones), S34Y (n=4 clones), S34F (n=4 clones) or S34F(TTC) (n=4 clones) amino acid substitutions in U2AF1, compared to parental A549 cells (n=4 clones) as determined by rMATS (v4.1.0)^63^. Detected splicing events consist of 5 categories: skipped exons (SE), retained introns (RI), mutually exclusive exons (MXE), alternative 5’ splice site usage (A5SS) and alternative 3’ splice site usage (A3SS).

**Extended Data Fig. 3:**
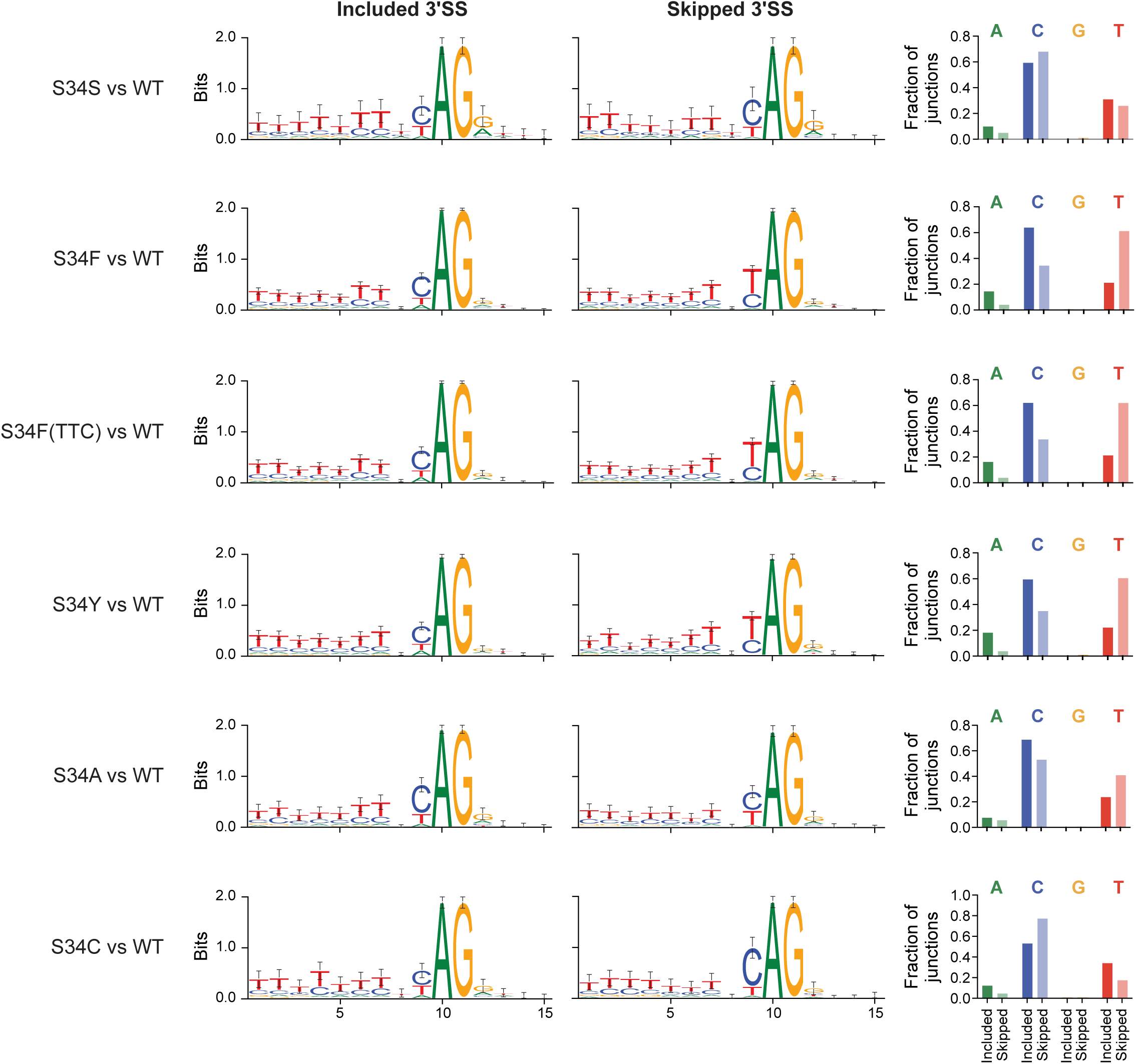
3’ splice site preferences for all engineered U2AF1 variants. Splice site preferences were determined by examining the sequence surrounding the 3’ splice site in significantly included exons compared to the alternative skipped exon. This was performed for S34S (n=81 splicing events), S34F (n=947 splicing events), S34F(TTC) (n=784 splicing events), S34Y (n=516 splicing events), S34A (n=198 splicing events), and S34C (n=132 splicing events) cells compared to parental A549 cells, and WebLogos^66^ were produced for each data set. The fraction of included (opaque bars) and skipped (partially translucent bars) junctions with a given base preceding the AG dinucleotide is displayed on the right.

**Extended Data Fig. 4:**
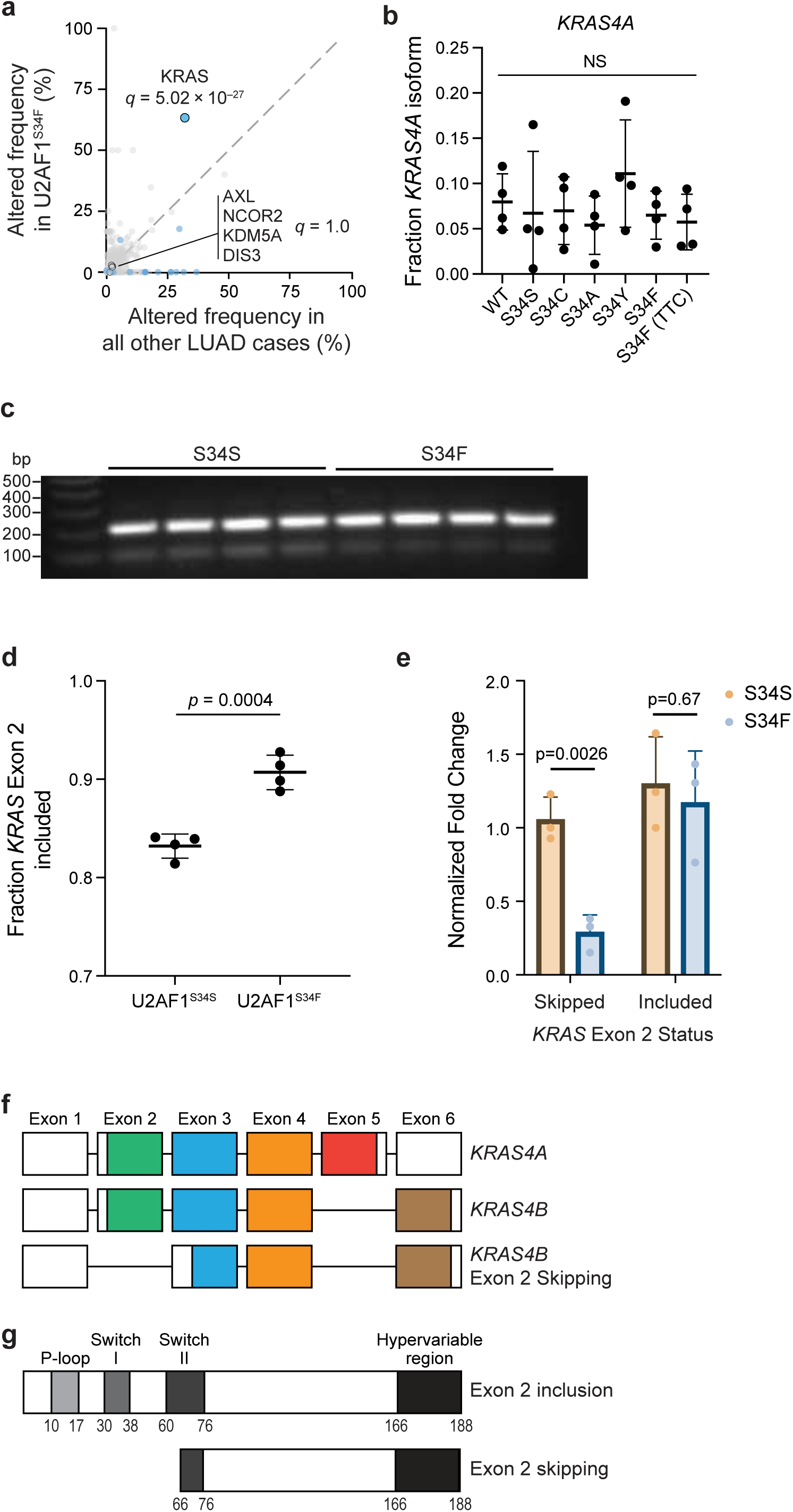
KRAS^G12S^-mutant cells undergo *KRAS* exon 2 skipping which can be reversed by U2AF1^S34F^ mutations. a) Frequency of genetic alterations in U2AF1^S34F^-mutant cancers compared to all lung adenocarcinomas as determined using the AACR Project GENIE dataset (v15.0)^11,29^. b) Quantification of the fraction of RNA-sequencing reads with *KRAS4A* isoform usage across parental A549 cells or those harboring S34S, S34C, S34A, S34Y, S34F or S34F(TTC) mutations (n=4 clones for each). Statistical analysis comparing each variant to parental A549 cells is shown (q ratio <1.02, DF=21, p>0.81 for all variants). c) RT-PCR detection of *KRAS* exon 2 skipping in A549 cells with U2AF1^S34S^ (n=4 clones) or U2AF1^S34F^ (n=4 clones) mutations. d) Quantification of the fraction of *KRAS* transcripts with exon 2 inclusion from c) (t=6.99, df=6, 95% confidence interval = 0.0487 to 0.101, p=0.0004). e) qRT-PCR quantification of *KRAS* exon 2 skipping or inclusion in A549 cells with U2AF1^S34S^ (n=3 clones) or U2AF1^S34F^ (n=3 clones) mutations. Fold change is normalized to Beta-actin expression. Statistical analysis comparing the amount of *KRAS* exon 2 skipping or inclusion in A549 cells with U2AF1^S34S^ or U2AF1^S34F^ mutations is shown (exon skipping: t=6.69, df=4, 95% confidence interval = -1.082 to -0.448, p=0.0026, exon inclusion: t=0.461, df=4, 95% confidence interval = -0.900 to 0.644, p=0.67). f) Diagram of *KRAS* gene structure showing exons used in *KRAS4A*, *KRAS4B*, and *KRAS4B* with exon 2 skipping. White segments represent untranslated regions. When exon 2 skipping occurs, this results in skipping of the translation start site and predicted use of an internal translation start site in exon 3. Diagram modified from Raso *et al.*^81^ g) Diagram of KRAS protein structure with and without exon 2 skipping. When exon 2 skipping occurs, translation is predicted to begin at an internal translation site at amino acid 66. Diagram modified from Kwan *et al.*^82^.

**Extended Data Fig. 5:**
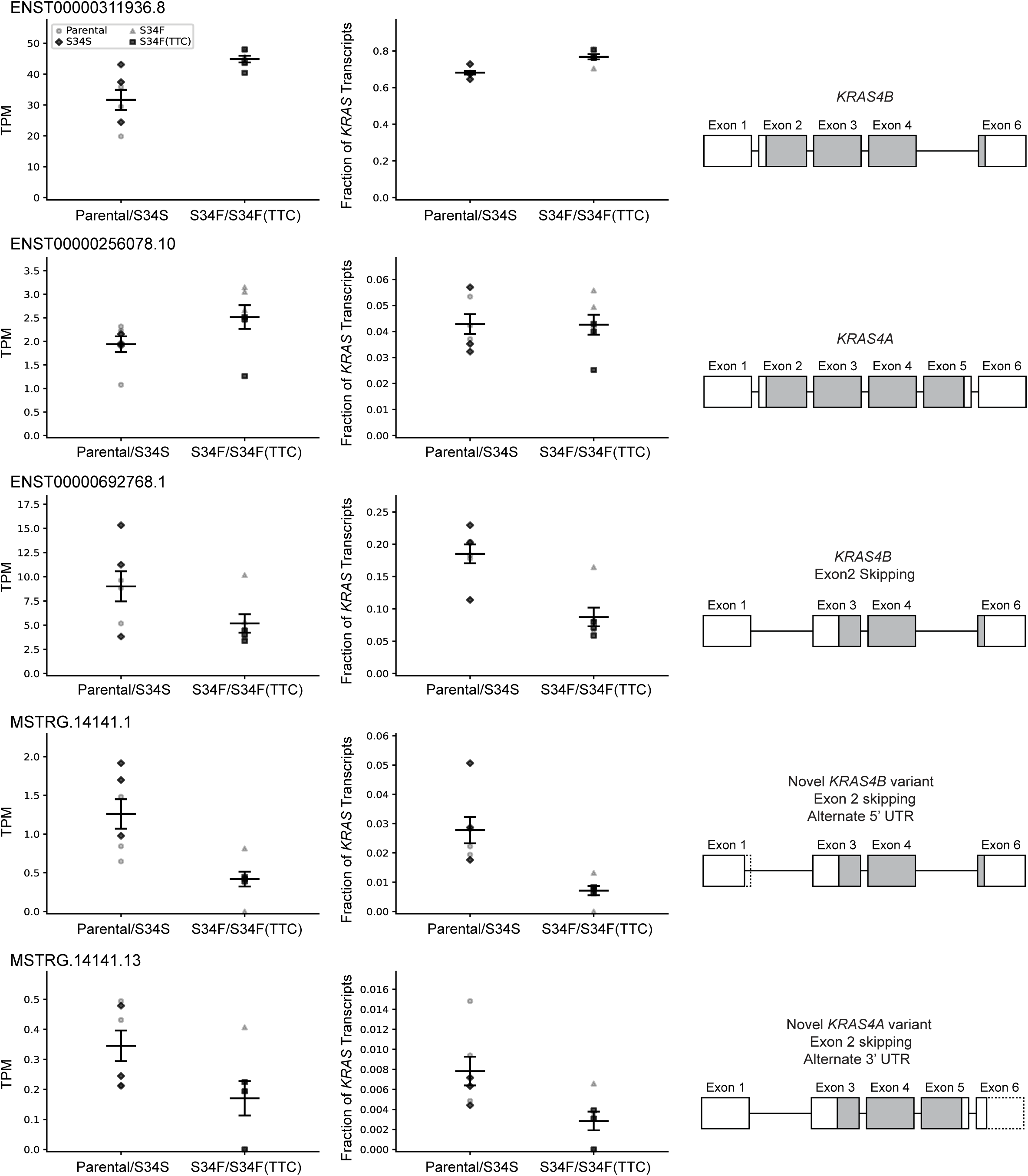
Long-read transcript usage for *KRAS4A, KRAS4B* and annotated *KRAS* exon 2 skipping transcripts. MSTRG14141.1 is a *KRAS4B* exon 2 skipping transcript observed using an alternative 5’ UTR. MSTRG14141.13 is a *KRAS4A* exon 2 skipping transcript observed using an alternative 3’ UTR. Transcripts per million (TPM) and fraction of *KRAS* transcripts are shown for each transcript. For each splicing diagram, boxes represent included exons, grey shading represents the predicted translation product, and dashed lines indicate alternative UTR usage.

**Extended Data Fig. 6:**
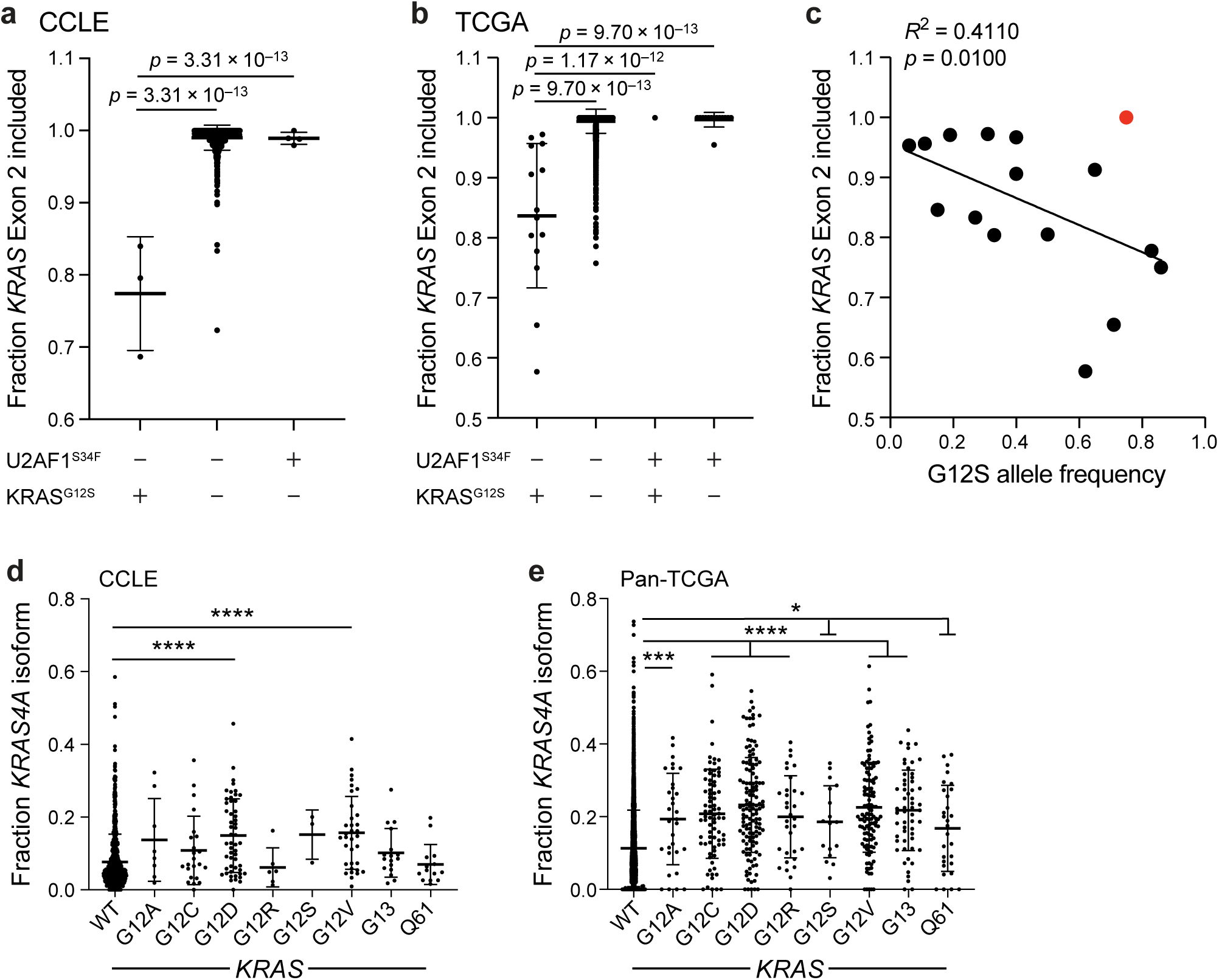
KRAS^G12S^-mutant cancer cell lines and patient tumors exhibit *KRAS* exon 2 skipping. a) Additional analysis of CCLE data from Fig. 1f, comparing *KRAS* exon 2 skipping in samples with KRAS^G12S^ and wildtype U2AF1 (n=3 cases), wildtype KRAS and wildtype U2AF1 (n=1001 cases), and wildtype KRAS and U2AF1^S34F^ (n=4 cases). Statistical comparison of wildtype U2AF1 + KRAS^G12S^ samples vs. wildtype U2AF1 + wildtype KRAS (q ratio=29.76, DF=1005, p=3.31x10^-13^) and comparison of wildtype U2AF1 + KRAS^G12S^ vs. U2AF1^S34F^ + wildtype KRAS (q ratio=22.44, DF=1005, p=3.31x10^-13^) are shown. b) Additional analysis of TCGA data from Fig. 1g, comparing *KRAS* exon 2 skipping in samples with KRAS^G12S^ and wildtype U2AF1 (n=14 cases), wildtype KRAS and wildtype U2AF1 (n=6028 cases), KRAS^G12S^ and U2AF1^S34F^ (n=1 case), and wildtype KRAS and U2AF1^S34F^ (n=14 cases). Statistical comparisons between wildtype U2AF1 + KRAS^G12S^ samples and wildtype U2AF1 + wildtype KRAS samples (q ratio=40.19, DF=6053, p=9.70x10^-13^), U2AF1^S34F^ + KRAS^G12S^ samples (q ratio=10.77, DF=6053, p=1.17x10^-12^) and U2AF1^S34F^ + wildtype KRAS samples (q ratio=28.92, DF=6053, p=9.70x10^-13^) are shown. c) Correlation between the fraction of *KRAS* exon 2 inclusion and variant allele frequency of KRAS^G12S^ in pan-cancer patients from The Cancer Genome Atlas dataset (n=16 cases, F=9.07, DFn=1, DFd=13, R^2^=0.411, p=0.0100)^24^. Red sample indicates a KRAS^G12S^-mutant sample which also contains a U2AF1^S34F^ mutation, and was excluded from statistical analyses. d) Quantification of the fraction of RNA-Sequencing reads with *KRAS4A* isoform usage for cell lines from (CCLE)^35^ with wildtype KRAS (n=836), KRAS^G12A^ (n=8), KRAS^G12C^ (n=23), KRAS^G12D^ (n=53), KRAS^G12R^ (n=6), KRAS^G12S^ (n=3), KRAS^G12V^ (n=34), KRAS^G13^ (n=17), or KRAS^Q61^ (n=14) mutations. Statistical comparisons between wildtype KRAS and KRAS^G12D^ (q ratio=6.48, DF=985, p=1.14x10^-9^), and between wildtype KRAS and KRAS^G12V^ (q ratio=5.79, DF=985, p=7.54x10^-8^) are shown. e) Quantification of the fraction of RNA-Sequencing reads with *KRAS4A* isoform usage for pan-cancer patient samples from (TCGA)^24^ comparing wildtype KRAS (n=7022) to KRAS^G12A^ (n=31, q ratio=4.21, DF=7509, p=0.0002), KRAS^G12C^ (n=79, q ratio=7.91, DF=7509, p<1x10^-15^), KRAS^G12D^ (n=131, q ratio=12.73, DF=7509, p<1x10^-15^), KRAS^G12R^ (n=32, q ratio=4.60, DF=7509, p=3.41x10^-5^), KRAS^G12S^ (n=17, q ratio=2.83, DF=7509, p=0.0364), KRAS^G12V^ (n=121, q ratio=11.58, DF=7509, p<1x10^-15^), KRAS^G13^ (n=56, q ratio=7.32, DF=7509, p<1x10^-15^), or KRAS^Q61^ (n=29, q ratio=2.77, DF=7509, p=0.0435) mutations.

**Extended Data Fig. 7:**
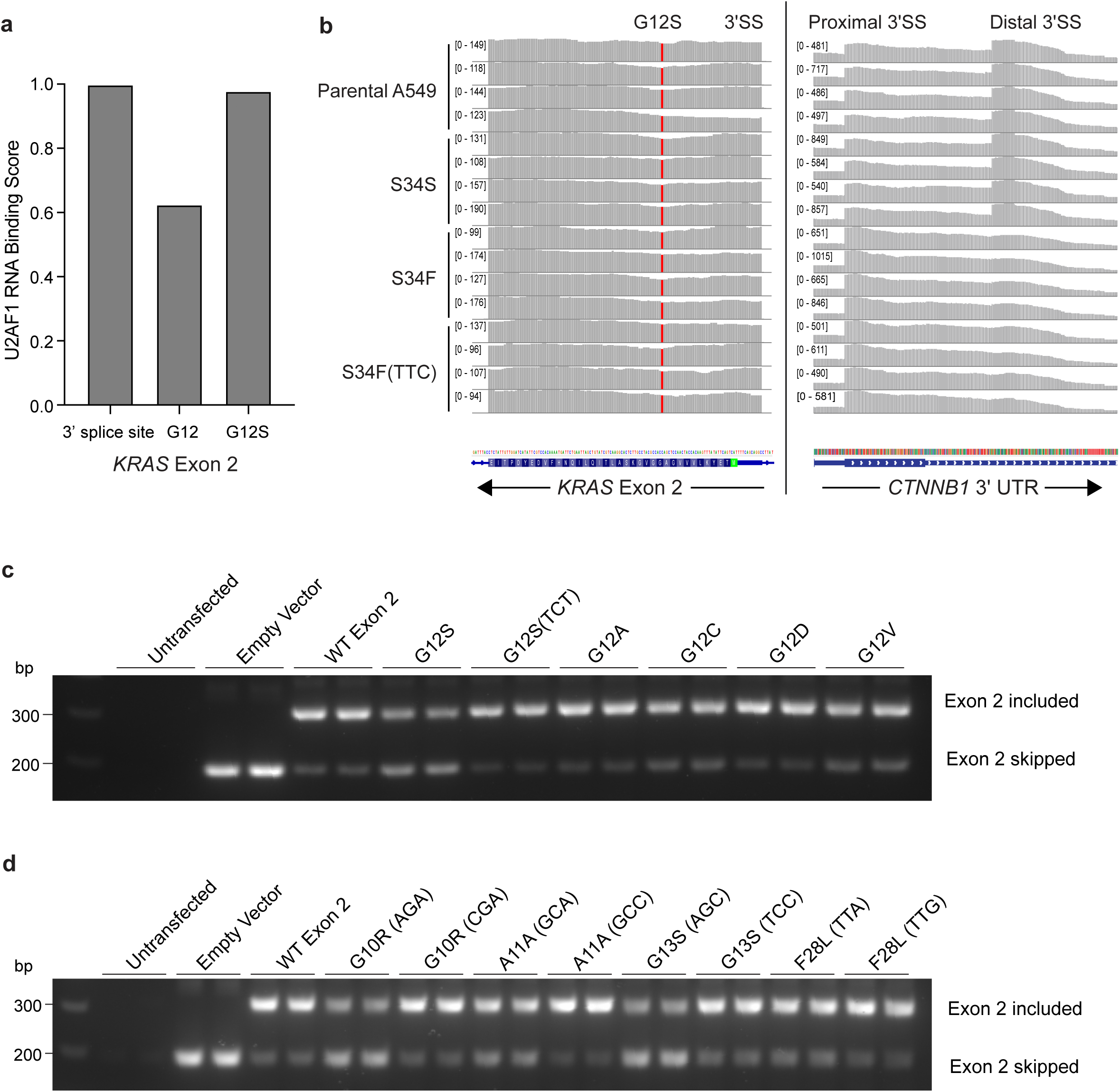
*KRAS* undergoes an exon 2 skipping event caused by the creation of a U2AF1 binding site upon KRAS^G12S^ mutation. a) U2AF1 binding score for an RNA sequence centered on the *KRAS* exon 2 3’ splice site, wildtype *KRAS* amino acid G12, or mutant *KRAS* G12S. Binding score was determined by RBPsuite (v1.0)^37^. b) Read count distribution across *KRAS* exon 2 or *CTNNB1* 3’UTR in parental A549 cells (n=4 clones) or those with U2AF1^S34S^ (n=4 clones), U2AF1^S34F^ (n=4 clones) or U2AF1^S34F(TTC)^ (n=4 clones) mutations. Reads were visualized using Integrative Genomics Viewer^64^. Red nucleotide in *KRAS* exon 2 indicates the homozygous KRAS^G12S^ mutation present in A549 cells. Arrow indicates the directionality of the gene. c) RT-PCR detection of *KRAS* exon 2 skipping for 293T cells transfected with *KRAS* exon 2 splicing reporter constructs. Cells were either untransfected (n=2), transfected with an empty vector without *KRAS* exon 2 (n=2), transfected with a wildtype *KRAS* exon 2 splicing reporter (n=2), or *KRAS* exon 2 with G12S (n=2), G12S(TCT) (n=2), G12A (n=2), G12C (n=2), G12D (n=2) or G12V (n=2) mutations. d) RT-PCR detection of *KRAS* exon 2 skipping for 293T cells transfected with *KRAS* exon 2 splicing reporter constructs. Cells were either untransfected (n=2), transfected with an empty vector without *KRAS* exon 2 (n=2), transfected with a wildtype *KRAS* exon 2 splicing reporter (n=2), or *KRAS* exon 2 with mutations introducing novel U2AF1 binding sites (AG dinucleotides) or synonymous mutations without U2AF1 binding sites. Mutations that generated novel AG dinucleotides include G10R(AGA) (n=2), A11A(GCA) (n=2), G13S(AGC) (n=2) and F28L(TTA) (n=2), while those that do not contain AG dinucleotides include G10R(CGA) (n=2), A11A(GCC) (n=2), G13S(TCC) (n=2), and F28L(TTG) (n=2).

**Extended Data Fig. 8:**
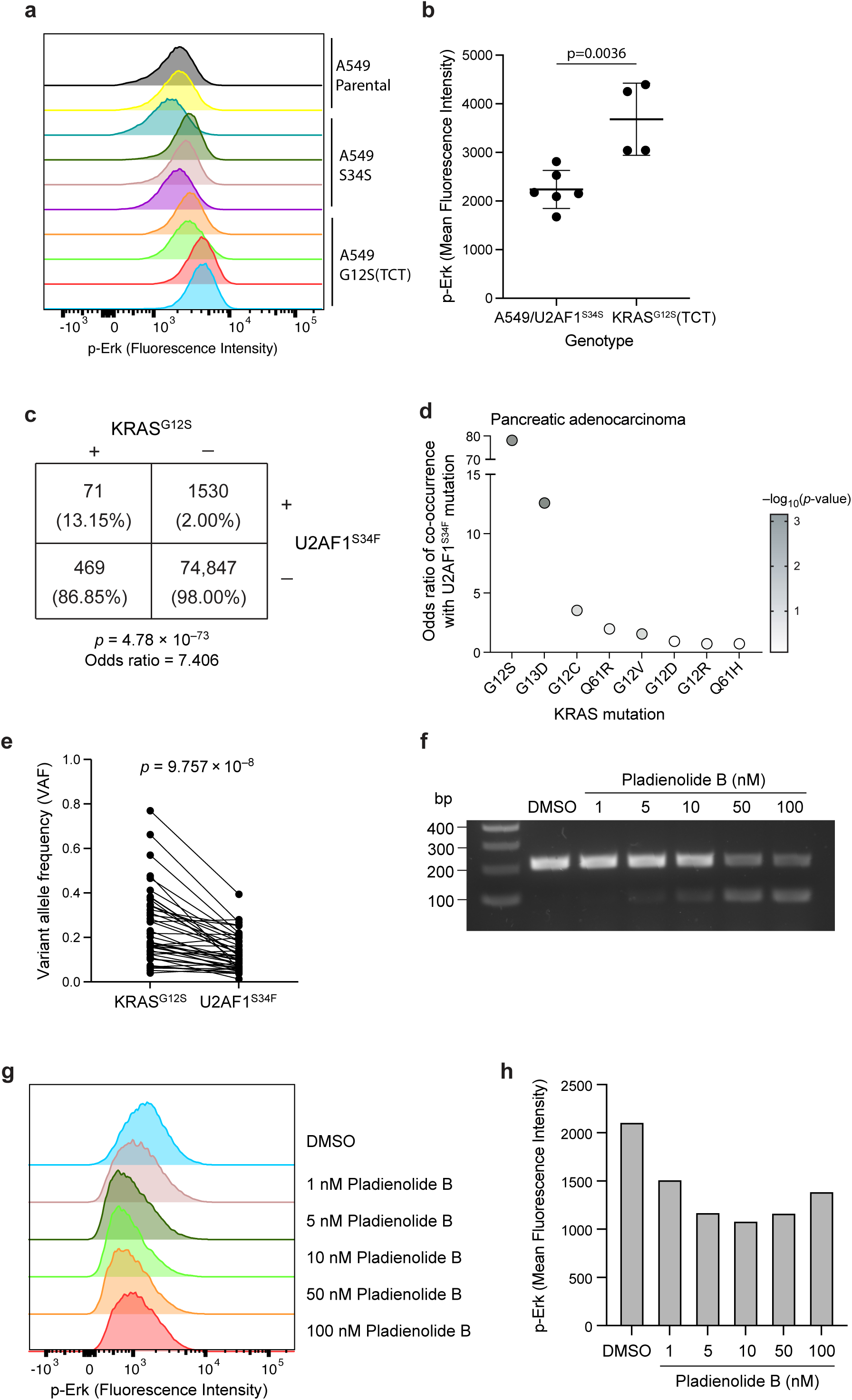
U2AF1^S34F^ mutations are enriched as secondary mutations in KRAS^G12S^- mutant cancers. a) Fluorescence intensity of p-Erk as measured by flow cytometry for parental A549 cells (n= 2 clones), U2AF1^S34S^-mutant A549 cells (n= 4 clones), or KRAS^G12S(TCT)^-mutant A549 cells (n= 4 clones). b) Quantification of the mean fluorescence intensity of p-Erk staining in parental A549 or U2AF1^S34S^ cells (n= 6 clones) or U2AF1^S34F^ cells (n= 4 clones) (t=4.07, df=8, 95% confidence interval = 627 to 2261, p=0.0036). c) Contingency plot analysis of U2AF1^S34F^ and KRAS^G12S^ mutations in lung adenocarcinoma patients from Foundation Medicine Inc. (n=62009 patients) and AACR Project GENIE (v14.0, n=14908 patients)^11^, including column percentages (odds ratio=7.406, p=4.78x10^-73^). d) Odds ratio of co-occurrence for U2AF1^S34F^ mutations and indicated KRAS mutations in pancreatic adenocarcinoma patients using data from AACR Project GENIE (v15.0, n=6528 patients)^11^. e) Paired analysis of the variant allele frequency of KRAS^G12S^ and U2AF1^S34F^ across 43 patient samples from the Foundation Medicine Inc. dataset containing both mutations and lacking CNVs at either locus (t=6.42, df=42, 95% confidence interval= -0.155 to -0.081, p= 9.757x10^-8^). f) RT-PCR detection of *KRAS* exon 2 skipping in NCI-H2023 cells treated with DMSO or 1 nM, 5 nM, 10 nM, 50 nM or 100 nM of the SF3B1 inhibitor Pladienolide B. g) Fluorescence intensity of p-Erk as measured by flow cytometry for NCI-H2023 cells treated with DMSO or 1 nM, 5 nM, 10 nM, 50 nM or 100 nM of Pladienolide B. h) Quantification of the mean fluorescence intensity of p-Erk staining in NCI-H2023 cells treated with DMSO or 1 nM, 5 nM, 10 nM, 50 nM or 100 nM of Pladienolide B.

**Extended Data Fig. 9:**
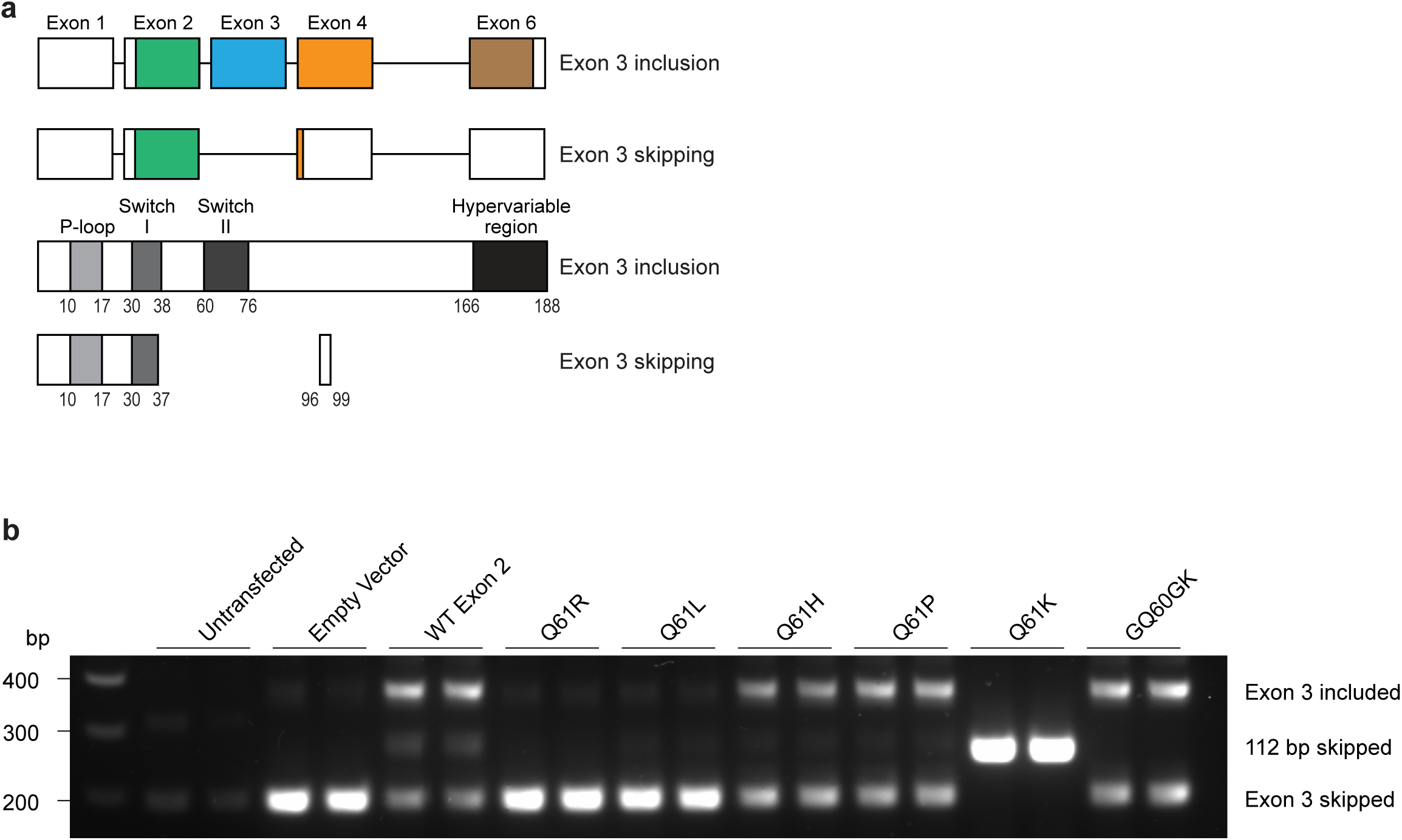
KRAS^Q61R/L^ mutations lead to *KRAS* exon 3 skipping. a) Diagram of *KRAS* gene and protein structure. Top shows exons used in *KRAS4B* with and without exon 3 skipping. White segments represent untranslated regions. When exon 3 skipping occurs, this results in early translation termination in exon 4. Bottom shows the KRAS protein structure with and without exon 3 skipping. When exon 3 skipping occurs, translation is predicted to terminate after amino acid 37, producing 3 incorrect amino acids before terminating. b) RT-PCR detection of *KRAS* exon 3 skipping for 293T cells transfected with *KRAS* exon 3 splicing reporter constructs. Cells were either untransfected (n=2), transfected with an empty vector without *KRAS* exon 3 (n=2), transfected with a wildtype *KRAS* exon 3 splicing reporter (n=2), or *KRAS* exon 3 with Q61R (n=2), Q61L (n=2), Q61H (n=2), Q61P (n=2), Q61K (n=2) or GQ60GK (n=2) mutations.

**Extended Data Fig. 10:**
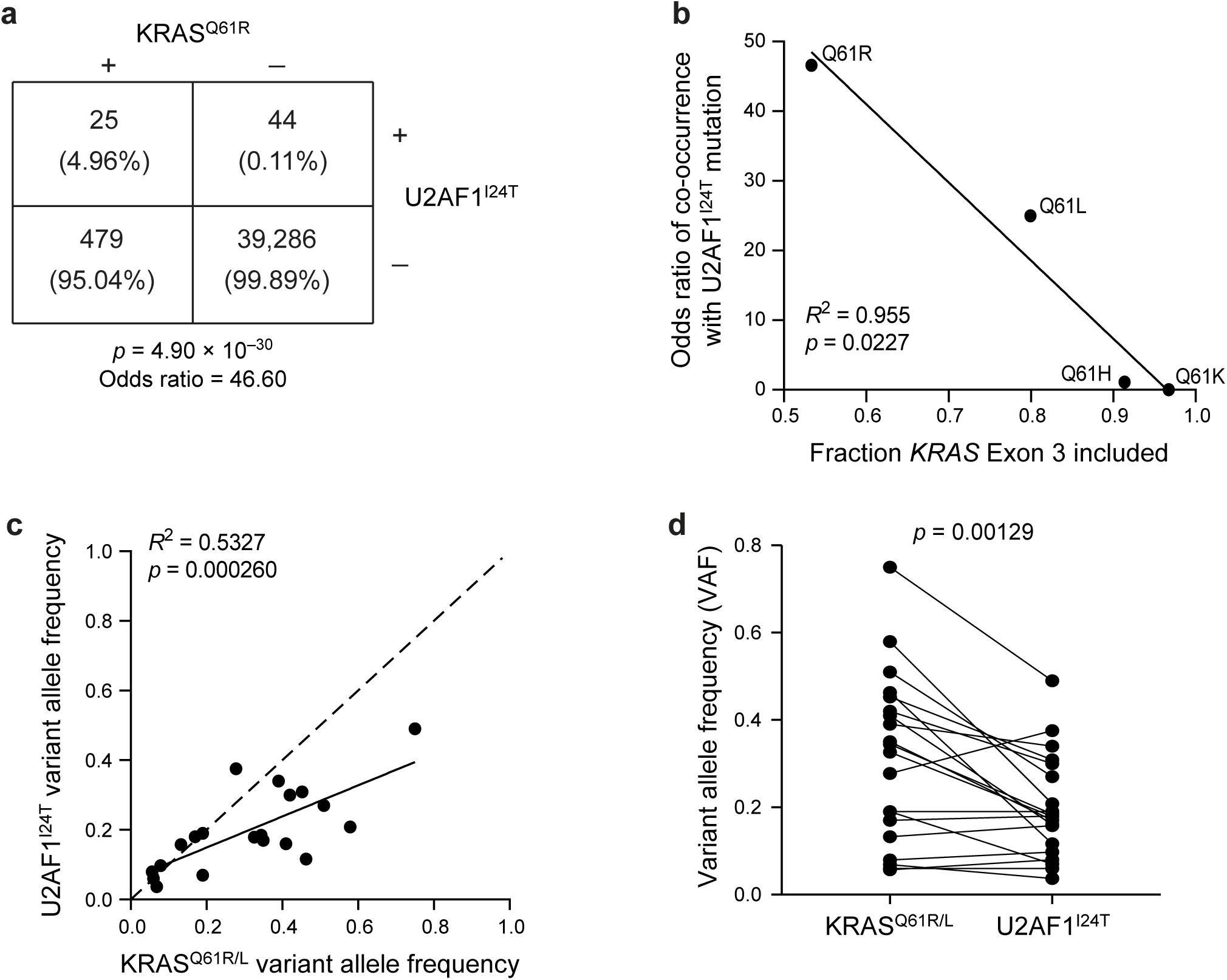
U2AF1^I24T^ mutations are enriched as secondary mutations in KRAS^Q61R/L^-mutant pancreatic cancers. a) Contingency plot analysis of U2AF1^I24T^ and KRAS^Q61R^ mutations in pancreatic cancer patients from Foundation Medicine Inc. (n=31,530 patients) and AACR Project GENIE (v16.1, n=8,304 patients)^11^, including column percentages (odds ratio=46.60, p=4.90x10^-30^). b) Odds ratio of co-occurrence of various KRAS^Q61^ mutations with U2AF1^I24T^ from Foundation Medicine Inc. (n=31,530 patients) and AACR Project GENIE (v16.1, n=8,304 patients)^11^ compared to the mean fraction of *KRAS* exon 3 inclusion for each Q61 mutation as quantified from TCGA^24^ (F=42.48, DFn=1, DFd=2, R^2^=0.955, p=0.0227). c) Correlation between the variant allele frequencies for KRAS^Q61R/L^ and U2AF1^I24T^ in 20 patient samples from the Foundation Medicine Inc. and AACR Project GENIE (v16.1) datasets containing both mutations and lacking CNVs at either locus (F=20.52, DFn=1, DFd=18, R^2^=0.5327, p=0.000260). Dashed line indicates a 1:1 ratio of KRAS^G12S^ and U2AF1^S34F^ allele frequencies. d) Paired analysis of the variant allele frequency of KRAS^Q61R/L^ and U2AF1^I24T^ across 20 patient samples from the Foundation Medicine Inc. and AACR Project GENIE (v16.1) datasets containing both mutations and lacking CNVs at either locus (t=3.77, df=19, 95% confidence interval=-0.175 to -0.050, p=0.00129).

## Notes

### Summary of Updates

New results added to Figure 2; new results added to Figure 3 and combined with previous Figure 4; new results added to new Figure 4; new Figure 5; new results added to Extended Data Figure 4, new Extended Data Figure 5; data from previous Extended Data Figure 4 split into new Extended Data Figure 6; new Extended Data Figure 7; new Extended Data Figure 8; new Extended Data Figure 9; results from previous Extended Data Figure 7 moved to new Extended Data Figure 10; updated author list, manuscript text, methods, references and figure legends.

